# KIMMDY: a biomolecular reaction emulator

**DOI:** 10.1101/2025.07.02.662624

**Authors:** Eric Hartmann, Jannik Buhr, Kai Riedmiller, Evgeni Ulanov, Boris N. Schüpp, Denis Kiesewetter, Daniel Sucerquia, Camilo Aponte-Santamaría, Frauke Gräter

## Abstract

Molecular simulations have become indispensable in biological research. Their accuracy continues to improve, but directly modelling biochemical reactions – central to all life processes – remains computationally challenging. Here, we present a biomolecular reaction emulator that models reactions across conformational ensembles using kinetic Monte Carlo. Our method, KIMMDY, is capable of handling dynamic, large-scale systems with successive, competing reactions, even on the second timescale or slower. It leverages graph neural networks for large-scale prediction of reaction rates, while also being capable of using simpler physics-based or heuristic models. We validate our approach against experimental data and showcase its power and versatility through a series of applications, including radical reactions, nucleophilic substitutions, and photodimerization. Example systems span proteins and DNA. KIMMDY aids the understanding of biochemical reaction cascades in complex systems, helps to re-interpret experimental data, and can inspire future wet-lab experiments.

## 1 Introduction

Biomolecules are inherently reactive. Their reactivity enables essential biological processes, Nature makes use of a plethora of ingenious biochemical reactions, from nucleophilic substitutions and additions to rearrangements and redox reactions to drive essential biological processes, including metabolism, signalling, and energy transfer. Beyond these tightly controlled and specific reactions, biomolecules undergo a wide range of unspecific reactions during their lifespan while they are subjected to varying conditions, being it pH, light, or oxidative stress, causing constant molecular damage under physiological conditions.

These biochemical processes happen within and are driven by the complex, crowded, and dynamic molecular environment inside and outside cells. Molecular simulations of such systems have given unprecedented insights into the inner workings of cells, thereby helping to overcome the fundamental limitation in the spatio-temporal resolution of experiments. Length scales and molecular complexities of the atomistic or coarse-grained models in biomolecular simulations are approaching, or already reaching, those of whole cells [1, 2]. However, these simulations lack the key feature of life: reactivity, which has to be sacrificed for efficiency. Molecular mechanics (MM) force fields are the prime choice to handle large system at sufficient time scales, but bonds can not break and form in these models. Monitoring the interplay of molecular motion and chemical events, however, is key to understanding and designing complex molecular systems.

To overcome the limitations of classical simulations, reactive force fields [3, 4], or schemes that allow for specific reactions to occur, such as the empirical valence bond method [5], or constant pH simulations [6] have been put forward. Also the more recent machine-learned potentials (MLPs), currently transforming the role of atomistic modelling, are by design reactive [7–12].

All of these models – MLPs, classical reactive, and even the highly efficient unreactive force fields – face the fundamental limitation that the reachable timescales are far below those of most biochemical reactions. The same naturally applies to hybrid approaches such as Quantum Mechanics/Molecular Mechanics (QM/MM) and Machine Learning/Molecular Mechanics (ML/MM), whose performance is kept back by the slower method of the two. Even if such hybrid approaches become more efficient in the future, they are unfeasible when it comes to the rich chemistry of biomolecules, being it post-translational modifications or radical chain reactions, where reactive regions can essentially span the whole system and timescales of reactions can reach seconds. Overcoming these limitations would enable large-scale simulations of reactive biomolecular systems.

We here present KIMMDY, a biomolecular reaction emulator (Fig. 1). KIMMDY searches for reactions within a dynamic molecular system and predicts reaction rates based on an emulator, that is, a machine-learned, heuristic or physical surrogate model.

**Fig. 1.**
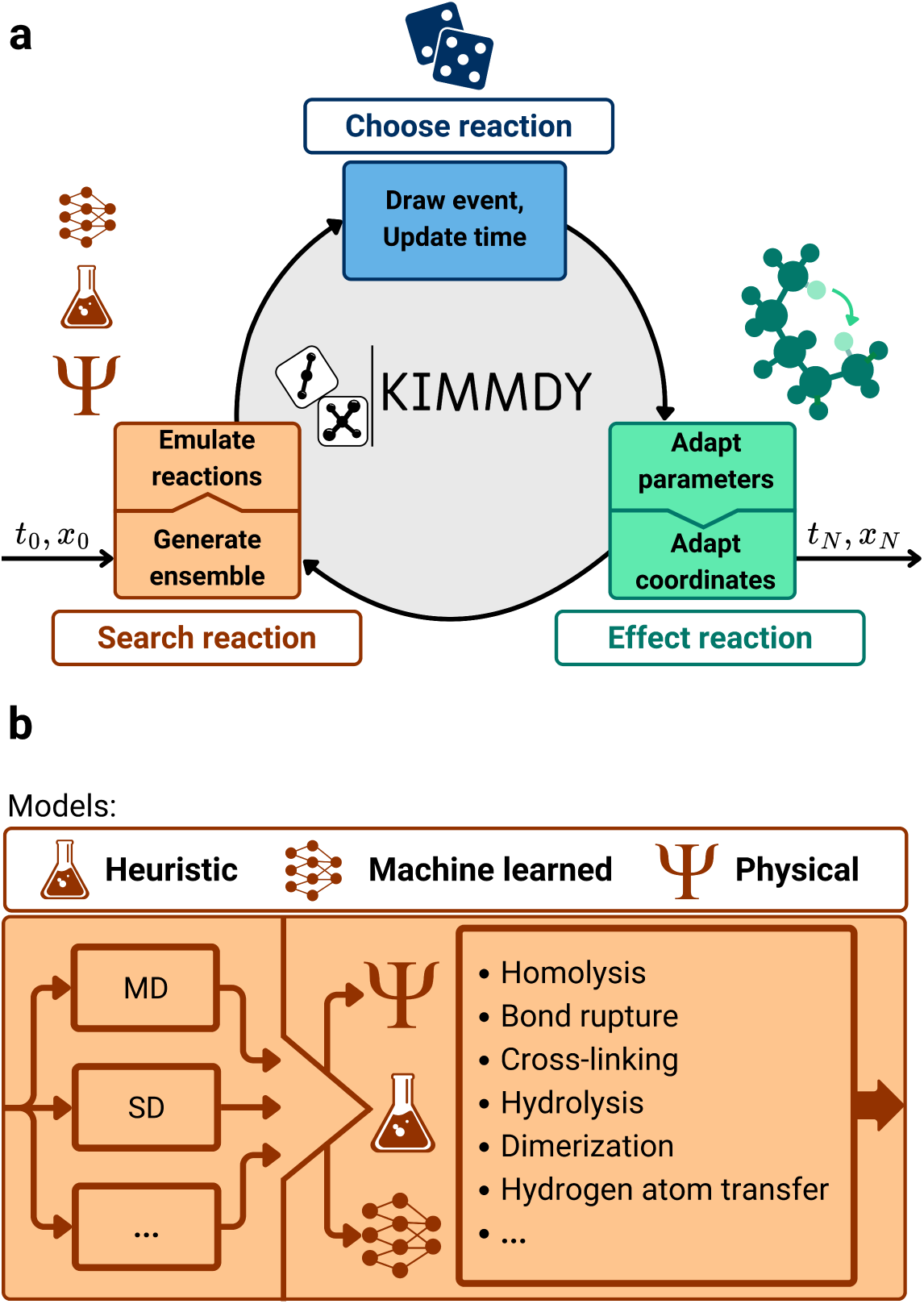
KIMMDY as a biomolecular reaction emulator. **a**, Visualization of the central workflow. KIMMDY searches for reactions within a dynamic molecular system, predicts reaction rates based on a learned, heuristic or physical model, chooses on of those reactions by kMC, and then effects the reaction. **b**, Detailed scheme of the “Search reaction” step of KIMMDY, displaying possible input ensemble generators and reaction types. MD and SD refer to Molecular and Stochastic Dynamics.

KIMMDY subsequently chooses one of those reactions in a kinetic Monte Carlo (kMC) step, effects this reaction and generates a new conformational ensemble to emulating further reactions (Fig. 1a). KIMMDY can predict reaction dynamics in highly reactive and large biomolecular systems, can cope with reaction cascades therein as well as competing reactions, and can in principle handle reactions at virtually any timescale. While combinations of MD and kMC have been used previously[13–15], our approach directly takes conformational ensembles into consideration and harnesses the power of graph neural networks (GNN) to rapidly predict ensemble-based reaction barriers [16]. At the same time, we profit from the validated accuracy and speed of classical biomolecular physics-based force fields [17, 18] and the ease of extending them to new chemistries by ML [19]. We validate KIMMDY against experiments and demonstrate its applicability to open and closed-shell reactions and protein and nucleic acid systems (Fig. 1b). KIMMDY is implemented as a modular platform that can be extended to arbitrary reactions, providing reaction rates and force fields are accessible via ML-models or through experimental data.

## 2 Results

### 2.1 Emulated reaction dynamics correctly predict radical reactions in small molecules

As a first area of application, we chose hydrogen atom transfer (HAT). HAT is ubiquitous in biomolecules and soft matter, can occur unspecifically and in cascades of many subsequent radical migration steps, often on timescales of seconds, and is thus a reaction type where standard approaches such as QM/MM or direct MD simulations with machine-learned potentials fail. We started with validating KIMMDY for HAT within small hydrocarbon radicals, for which experimental results are available. The systems consisted of n-alkane derived radicals, starting with the 1-propyl radical up to the 1-octyl radical (Fig. 2a). The machine-learned barrier prediction model used to infer rates for kMC was trained on barriers calculated at the DFT level for HAT reactions for proteins [16], which we hypothesized to sufficiently generalize to hydrocarbons (see Supplementary Methods).

**Fig. 2.**
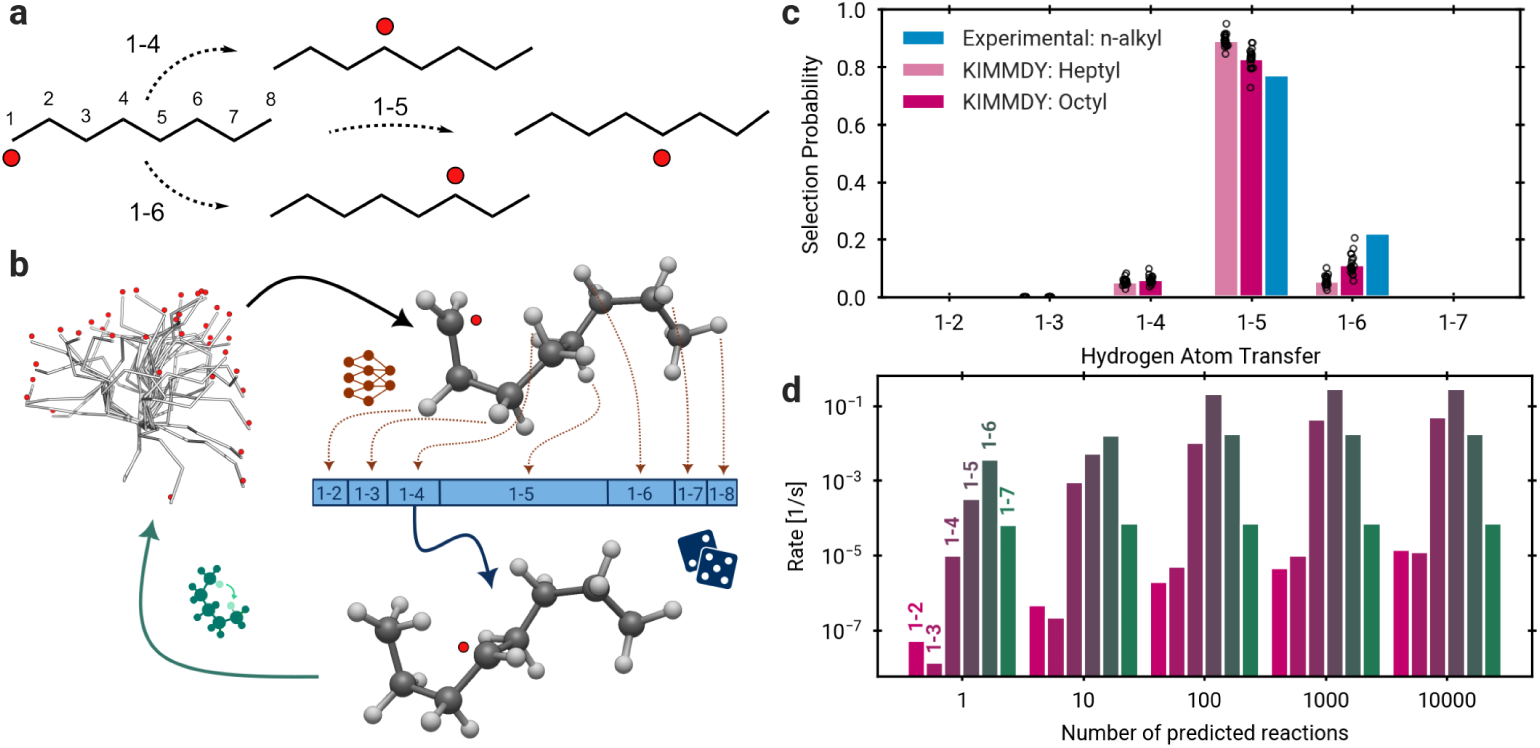
Validating KIMMDY: Hydrogen atom transfer (HAT) in n-alkyl radicals. **a**, Illustration of 1-4, 1-5 and 1-6 HATs in an 1-octyl radical. **b**, KIMMDY HAT prediction cycle. KIMMDY searches for possible HAT reactions within an MD-generated conformational ensemble, infers their barriers from an emulator, chooses one reaction by kMC, adjusts the topology, and a new MD simulation commences. **c**, Probability of HAT reactions as derived with KIMMDY for heptyl and octyl radicals, and compared with the experimental probability of n-alkyl (see Supplementary Methods). Circles indicate a single 100ns KIMMDY run and coloured bars signify the mean. For shifts that were not available experimentally, i.e. 1-2 and 1-7, the rates are shown in Fig. A1a, but the probabilities are set to 0 in c. Hydrogen shifts between primary carbons can not be detected in experiments and are excluded here. **d**, Mean rates for different number of predicted reactions (averaged over 4 runs, 10ns). Further details in Supplementary Notes.

Figure 2b shows the typical KIMMDY workflow, exemplified with an octyl radical. Figure 2c shows the selection probability for different HAT reactions at 500 K. We chose the mean of the experimentally-derived rates as reference for a given hydrogen shift divided by the sum of all mean experimentally-derived rates of all considered shifts. Similarly, we extracted the selection probability for these shifts from KIMMDY simulations. In line with the experimental results, our method correctly identifies 1-5 as the dominant HAT, due to a favourable 6-ring configuration of the transition state, and also accurately predicts the overall trend of the 1-n shift probabilities, particularly, the low probabilities of transitions with high ring strain energy of the transition state (1-2 and 1-3) [20]. The mean rate for each shift quickly converges for increasing number of barrier predictions per run (Fig. 2d, see the Supplementary Notes for further discussion). Thus, KIMMDY robustly reproduces the overall reaction dynamics, with only slight deviations for the most prominent shifts relative to each other.

The overall correct ranking of HAT reactions is also evident when directly comparing absolute reaction rates (Fig. A1 a). However, the GNN-based emulator underestimates the absolute rates. A reason for this could be the training data of the GNN-based emulator, which included constrained transition states for which only directly involved atoms were optimized [16]. Additionally, tunnelling is ignored, which is known to increase rates by up to two to three magnitudes at 300 K [21]. Including zero-point vibrational energy corrections in the training data set of the emulator as well as fine-tuning of the model to alkyl radicals might further improve the results. Notwithstanding this underestimation, KIMMDY accurately reproduces the expected reaction probabilities, the decisive quantity in the application scenario of our reaction emulator. Also, as demonstrated by our validation, KIMMDY with learned barriers can be readily applied even to systems (n-alkanes) not explicitly included into the training data set (proteins).

### 2.2 KIMMDY yields novel reaction pathways for protein radicals

Having validated our reaction emulator for HAT, we move on to the first application in a biological system. Here, radical species are the product of homolytic cleavage and subsequently migrate through the system in several consecutive HAT reactions (Fig. 3a).

**Fig. 3.**
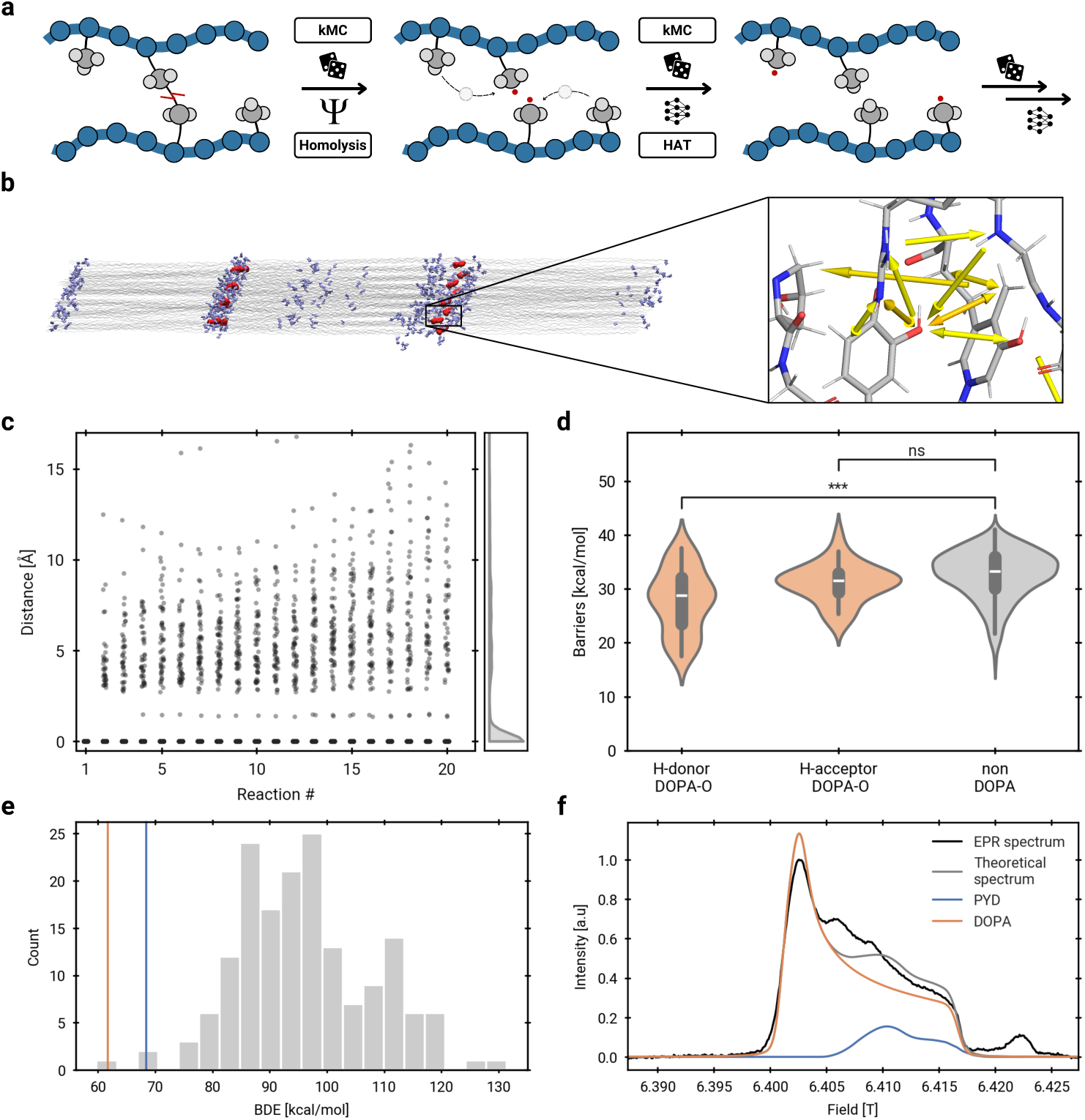
DOPA and PYD scavenge homolysis-derived radicals. **a**, Consecutive reactions of radicals in KIMMDY initiated with homolytic cleavage and followed by multiple HATs. Unpaired electrons are shown as red dots. **b**, Structure of collagen fibril with backbone atoms (silver), potential DOPA sites (light blue) and PYD crosslinks (dark red). The zoom shows HATs as yellow arrows at a crosslink site. **c**, Distance of radical atoms to the homolytically cleaved PYD C*α* and C*β* atoms for 36 simulations with 8 radical atoms each over a maximum of 20 consecutive HAT reactions. The density is shown for before the 20th reaction. **d**, Distribution of HAT barriers for reactions involving the DOPA hydroxy group as H-donor (n=49, median=28.7 kcal/mol), H-acceptor (n=41, median=31.4 kcal/mol) or no DOPA (n=467, median=33.2 kcal/mol). The differences between H-donor DOPA hydroxy group and non-DOPA are significant using Welch’s t-test (p=3.9e-7). **e**, Bond dissociation energy BDE of PYD (blue line) and DOPA (orange line) in a distribution of BDE of abstractions processes in different amino acids (from [25], gray bars). **f**, Experimental EPR spectrum of rat tail collagen (black), reproduced with permission from Kurth *et al.* [23] (black) is compared with the spectrum of DOPA (orange), previously proposed as a key contributor to the signal. The spectrum of the PYD crosslink (blue), a candidate proposed by KIMMDY, can explain unresolved parts of the experimental data.

Collagen, formed from aligned and crosslinked triple helices (Fig. 3b), harbours radicals from homolytic cleavage when subjected to mechanical stress [22]. Migration of these mechanoradicals to DOPA, a post-translationally modified catechol-containing amino acid, was proposed based on thermodynamic arguments and electron paramagnetic resonance (EPR) data [22, 23]. While homolysis sites [24] and the role of DOPA as a radical scavenger [23] have been extensively studied, KIMMDY offers the unique capability to scrutinize radical transfer pathways from homolysis sites to radical scavengers.

In a collagen fibril system of 2.6 million atoms, we observe a plethora of radical transfer pathways within 600 emulated HAT reactions. Through HATs and conformational changes, radicals move tens of Angstroms away from the homolytically cleaved atoms (Fig. 3c). A wide range of amino acids host radical species during the simulations, and side chain radicals were more frequent than backbone radicals (Fig. A2a,b). HATs are mainly intermolecular or with more than six bonds in between H-donors and H-acceptors and only include few 1-2, 1-3, or 1-4 transfers, consistent with the above simulations of alkanes (Fig. A2c). As expected for a radical scavenger, barriers are significantly lower for reactions with a DOPA hydroxy group H-donor compared to those not involving DOPA (Fig. 3d). The median barrier difference is 4.5 kcal/mol, which, using the Eyring equation (Supplementary Notes), amounts to 1800 times faster reactions. We also sampled direct HATs from the homolytic cleavage site to the hydroxy group of DOPA (Fig. A2b). Thus, DOPA radical species are kinetically accessible in collagen under stress. We do not observe the DOPA hydroxy radical to be less reactive as hydrogen acceptor compared to other protein radicals, and attribute this to rapid HATs between multiple adjacent radical scavengers (Fig. A2e,f).

Interestingly, a large population of radicals resulting from crosslink cleavage (C*α* and C*β* atoms of the trivalent pyridinium-crosslink PYD) do not migrate during the reactive simulation (Fig. 3c). This indicates that they are relatively stable radicals, compared to more reactive H-acceptors formed in subsequent reactions which often migrate further. In a multiple hypothesis testing scenario, we tested for amino acid H-donors with a significantly lowered barrier (Fig. A2d). Here, the hydroxy-group of the broken PYD is identified as a frequent H-donor. This moiety has not been investigated as potential radical scavenger before, but is ideally positioned and kinetically accessible according to KIMMDY.

We asked if PYD can also thermodynamically serve as radical sink in collagen by comparing its bond dissociation energy (BDE) for hydrogen abstraction to those in all twenty amino acids and DOPA [25] (Fig. 3e). Indeed, PYD and DOPA have the lowest BDEs, which implies that both are exceptionally good radical scavengers. A previously measured EPR absorption spectrum of stretched collagen [23] also supports the notion of PYD as a plausible radical candidate. It supports DOPA as the major radical species, but exhibits an otherwise unexplained region, which aligns with the computed PYD spectrum (see Supplementary Methods). Including PYD improves the overall fit to the experimental data, increasing the *R*^2^ from 0.89 to 0.98 (Fig. 3f and Fig. A3). Thus, KIMMDY lets us re-interpret experimental data and identify an overlooked radical-stabilizing moiety in collagen.

### 2.3 KIMMDY can model competing reactions in biologically relevant systems

In most biochemical systems, different reactions can directly compete with each other, leading to vastly different products. To showcase the ability of KIMMDY to deal with different reaction types, we compare homolytic and heterolytic bond cleavage in proteins. While homolytic cleavage leads to subsequent chain reactions and detectable radicals (see Sec. 2.2), heterolytic cleavage is a closed-shell reaction and in case of proteins involves the attack of water, i.e. is a hydrolysis reaction (Fig. 4A). Both reactions can be promoted by force, both have been observed experimentally [22, 24, 26, 27], and their competition depends on environmental effects, such as solvent accessibility or pH.

**Fig. 4.**
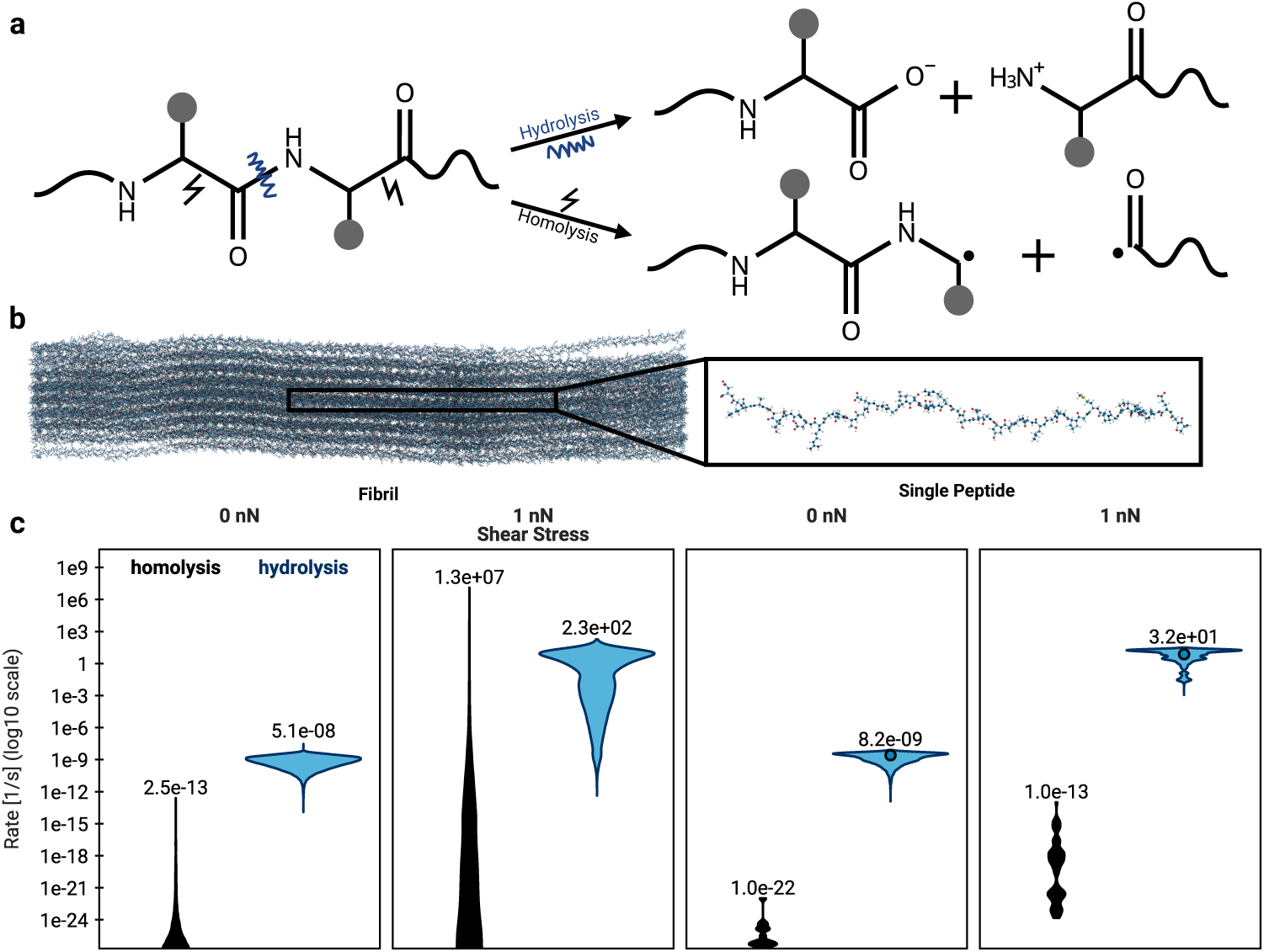
Hydrolysis vs. Homolysis. **a**, Schematic representation of a peptide that undergoes either base-catalysed hydrolysis or homolytic cleavage. **b**, Molecular render of a collagen fibril and a single peptide. **c**, Comparison of the reaction rates for two competing reactions, homolysis and hydrolysis based on a physical model for homolysis [29] and a heuristic model [26] for hydrolysis. The value of the highest rate for each setup is shown above the respective distribution. The circles in the hydrolysis rate distributions of the single peptide denote the experimental reference value for the respective force.

Direct comparisons between the two reactions using combined quantum mechanical and molecular mechanical simulations (QM/MM) are possible, but fundamentally limited by sampling and computational cost. This is where KIMMDY’s modular system comes into play, providing an emulator for their competition in a realistic biological setting. The reaction rates for homolysis are modelled according to a physics-based Bell model [28, 29] and those for hydrolysis via a heuristic model based on force-clamp experiments [26], pH, and surface accessibility, which demonstrates the ease of using KIMMDY even when conformation-dependent learned rates are not at hand.

As expected, in the absence of an external pulling force, both hydrolysis and homolysis yield low rates, that is, spontaneous fragmentation of peptides is unlikely at ambient conditions. When the protein systems are pulled at 1 nN, hydrolysis out-competes homolysis for a single peptide chain. However, for a densely packed and crosslinked collagen system, with all else being equal, homolysis is accelerated drastically and reaches rates of similar orders of magnitude as hydrolysis and outcompeting it when comparing the highest rates. This is due to areas of high stress concentration in the large collagen fibril system that push the reaction rates into a region of the force response curve where hydrolysis already tapers off and homolysis still scales (See Fig. A4b and [29]). This result is robust with regard to the model choice for hydrolysis (physical QM-based versus heuristic experiment-based) and other model parameters (see Supplementary Methods and Fig. A4).

Our results suggest that reactions that exhibit disparate scales of rates in simple molecules can become competitive in more complex molecular systems, with KIMMDY allowing direct conclusions on potential reaction outcomes, such as radical formation in the case of dense protein networks such as collagen.

### 2.4 KIMMDY reveals unexpectedly low quantum yields in DNA origami motifs

Upon UV irradiation, pyrimidine bases in DNA can form cyclobutane pyrimidine dimers (CPDs), which play a crucial role in the development of skin cancer [30]. In DNA nanotechnology, CPDs are used as covalent crosslinks between staples, enhancing the chemical stability of DNA origami structures [31]. MD studies have suggested that the frequency of DNA conformations favourable for dimerisation can be directly related to the quantum yields *ϕ* observed in experiments [32–34], with KIMMDY now offering a direct test of this scenario.

A distance and angle-based heuristic model, with parameters tuned to reproduce experimental quantum yields for small benchmark systems, was used to calculate reaction rates through KIMMDY (see Supplementary Methods). Simulation frames with rates above a defined threshold are counted as dimerisable configurations contributing to the quantum yield.

In the thymine dinucleotide (TdT) system, KIMMDY predicts two products, the expected *cis-syn* but also the *trans-syn* CPD isomer, albeit at much lower quantum yields (Fig. 5a and Fig. A5). Inspection of the underlying conformation shows that prior to the reaction, the thymine bases can undergo a *syn-anti* transition by rotation around the N-glycosidic bond (Fig. A5a,b). In experiments, the *cis-syn* isomer forms eight times more frequently than the *trans-syn* isomer during TdT irradiation [35]. Our KIMMDY results put forward a lower quantum yield, together with the *syn*-*anti* precursor state being less frequently populated, as explanation.

**Fig. 5.**
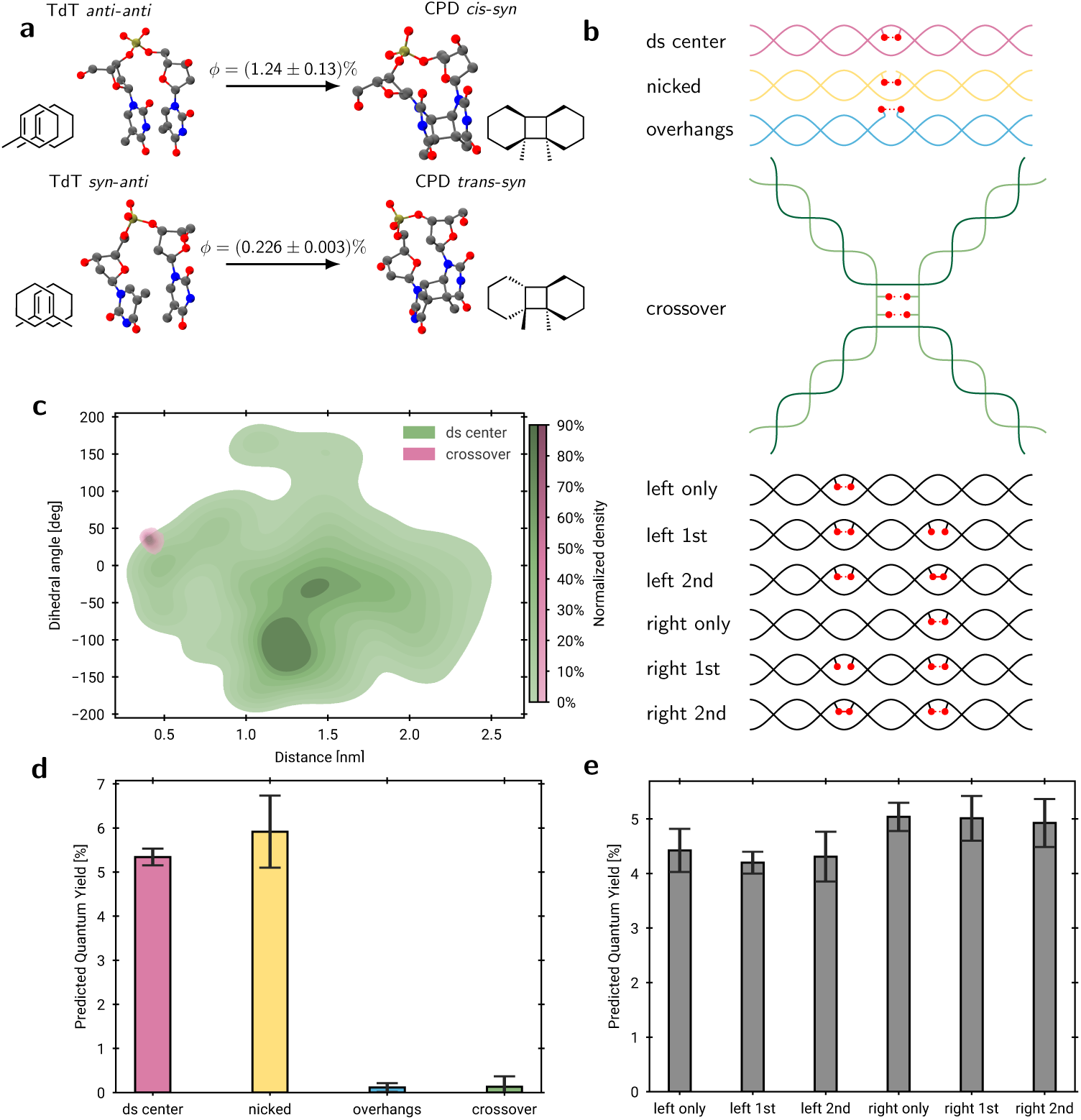
DNA systems investigated with KIMMDY. **a**, Reaction scheme of thymine-thymine dimerisation starting from the *anti-anti* -conformation or the *syn-anti* -conformation yielding *cis-syn* and *trans-syn* diastereomers with different predicted quantum yield *ϕ*. **b**, Structural schemes of tested DNA motifs. Red circles denote thymines. Solid connections between thymines indicate a dimer, dashed connections between thymines denote a reactive site considered by KIMMDY. **c**, Kernel density estimate (KDE) of distance and angle distributions of ds center and crossover systems determined from 10000 snapshots randomly sampled from three MD simulations (see Supplementary Methods), respectively. The KDE is plotted over the distance between reactive thymine double bonds and the dihedral angle between them. **d**, Predicted quantum yields for motifs commonly found in DNA origami (*n* = 3 for ds center, nicked, and overhangs, and *n* = 6 for crossover from separate 100 ns MD simulations). **e**, Predicted quantum yields for consecutive dimerisation systems (and respective control systems), in which the two reactive sites are five base pairs apart (*n* = 3 for left only, left 1st, right only, right 1st, *n* = 4 for right 2nd, and *n* = 5 for left 2nd from separate 100 ns MD simulations).

KIMMDY predicts quantum yields for overhang and crossover structures (Fig. 5b) that are substantially lower than for double-stranded (ds) or nicked motifs (Fig. 5d), despite overhangs and crossovers being commonly used to stabilize DNA origami *via* CPD crosslinking. This discrepancy may be explained by three reasons: long irradiation times used for origami stabilisation could compensate for the inherently low quantum yields; the distance and dihedral angle distributions show (Fig. 5c) an increased flexibility of the crossover system, which might be more restricted within the full origami structure; other photoproducts such as *cis-anti*, *trans-anti*, and 6-4 photoproducts (see Supplementary Methods), which are not accounted for in the current KIMMDY plugin, might be formed. We therefore propose that future experimental studies should investigate both the quantum yields and the diversity of photoproducts in DNA origami.

In systems with consecutive dimerisation sites (Fig. 5b), we observe no change in predicted quantum yields (Fig. 5e). This indicates that a close-by CPD separated by five base pairs from a second reactive site does not significantly affect the conformational space available for an additional dimerisation. This insight suggests that DNA previously damaged by CPD formation is not necessarily more susceptible to additional dimerisation, which may have implications for the formation of mutagenic lesions.

## 3 Discussion and Conclusion

KIMMDY offers the emulation of single reactions, reaction cascades, and competing reactions throughout molecular systems at a computational effort which is orders of magnitudes smaller than alternative approaches, being it at the ab initio, hybrid QM/MM, reactive force field, or MLP levels. Direct simulations with the latter and variations thereof are prohibitively slow for most biochemically relevant processes, as they are currently limited to the microsecond (or lower) timescale and require enhanced sampling for selected reactions.

The novelty of our work lies in harnessing the power of recently developed graph neural networks for two new tasks, (i) for emulating reactions and their barriers in a conformation-aware manner using a hybrid MD/ML approach and (ii) for learning a molecular mechanics force field as required for the physics-based ensemble generation in a reactive setting. We validated our reaction emulator by comparison to experimental and DFT data.

In our three exemplary applications, KIMMDY yields unexpected predictions to reinterpret former and inspire new experiments. First, we identify a new efficient radical stabilizing moiety in collagen, which can explain hitherto unexplained spectroscopic data and is relevant for our understanding of tissue mechanochemistry and ageing. Secondly, KIMMDY shows that the competition between open and closed shell biochemistry can substantially shift when going from simple to more complex structural environments, reconciling apparent contradictions of previous observations. Thirdly, KIMMDY predicts quantum yields of photoinduced dimerisation in DNA to be unexpectedly low in common DNA origami motifs and to be independent of previous nearby dimerisations — testable findings that are highly relevant to biomedical applications. Although KIMMDY is able to simulate a diverse set of systems and reactions, there are inherent limitations. KIMMDY reactions are selected and implemented for specific reaction types, thus it can not predict reaction types not anticipated by the researcher. For example, dimerisation could be extended to account for all possible photoproducts. Exploring reaction networks more widely[36] prior to KIMMDY can lift this limitation in the future. As shown in the n-alkyl case, HAT rates did not take tunnelling into account. Even though the probability of choosing a reaction is accurate, the time to the next reaction is off by a factor, limiting the interpretation of reaction times and their use when it comes to competing reactions. KIMMDY, however, offers the incorporation of such as tunnelling or other (quantum) effects into the emulator, which is subject of future work.

Heuristic models make the implementation of new reaction types straightforward, but their predictive power is limited. For example, dimerisation rate estimations based on transition states on the excited-state energy surface are expected to be more accurate and learnable [37]. The hydrolysis model uses a linear relation to the solvent-accessible surface area, again an assumption that an ML-based emulator can lift. Still, KIMMDY is applicable to problems for which efficient and sufficiently accurate heuristic models can be identified.

To conclude, KIMMDY is a versatile and highly extensible computational method to explore biochemical reactivity amidst the fluctuations of a dynamic molecular system. Large-scale simulations with KIMMDY include biochemical reactivity at little extra costs. We foresee KIMMDY to deliver new insights at many frontiers, from how enzymes work under tight regulation within the cell, to how biomolecules are permanently modified, damaged, degraded or repaired.

## 4 Methods

### 4.1 Adaptive kinetic Monte Carlo

KIMMDY simulates the time evolution [38] of reactive biopolymer species. It uses the rejection-free kMC (rf-kMC) algorithm[39, 40], which consists of the following steps: 1. From an initial state, create an event list of all possible transitions, 2. Draw a uniform random number u_1_ and select the event for which *F* (*p_i−_*_1_) *< u*_1_ *≤ F* (*p_i_*), where *F* is the cumulative function and *p_i_* the probability of event *i*. 3. Draw a uniform random number u_2_ to update the time according to 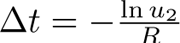, where R is the sum of all rates. 4. Carry out the event *i*. In KIMMDY, the simulated events are chemical reactions and transition rates are reaction rates.

Adaptive kMC [13] is a variation of rf-kMC where the event list for a state is calculated only if it is populated during the kMC simulation. This approach is beneficial if the state space is too vast or the number of possible events per state too large to precompute the event probabilities. In biopolymers and in general soft matter systems, most atoms exhibit unique reactivities because they are embedded in a certain structural context with electrostatics, solvent accessibility and steric effects. This dependence on the environment necessitates to calculate reaction rates individually for every set of reactant atoms, rendering their reactive simulation an ideal application for adaptive kMC.

In KIMMDY, a kMC step is divided into three tasks (Fig. 1a, see Supplementary Methods). First, possible reactions are sampled by generating a conformational ensemble of the current state, which then is used to predict reaction rates to neighbouring states either by a heuristic, physical or machine learned model (Fig. 1b). The second task comprises selecting the event and a corresponding update time according to the kMC algorithm. Finally, MD simulation parameters and coordinates are adapted according to obtain the product state. Several kMC steps can be chained in a cyclic process. We built KIMMDY to be modular, flexible and extensible.

### 4.2 Sample reaction

For reaction sampling, a conformational ensemble serves as the input to the reaction emulator, which predicts reaction rates for transitions to chemical states different from the current ensemble. In the applications presented here, a molecular system described by coordinates and simulation parameters is simulated using Molecular (MD) or Stochastic Dynamics (SD), but any other ensemble generator can be used, *e.g.* a learned one [41].

One approach to calculate the event list from the conformational ensemble generated with MD or by other means is by using ensemble averages of properties and relate them to reaction rates by physics-based or empirical models. Alternatively, we use a machine-learned model to predict transition rates from individual snapshots to calculate a constant average rate per reaction over the whole ensemble (see Supplementary Notes). This has the additional benefit of accounting for entropic effects by sampling how frequent highly reactive conformations are visited.

For the HAT application, using the conformation ensemble instead of a single structure representing a state, the prediction task is significantly simplified from emulating a complete transition state search to a local optimization of the reacting atoms. Still, the MD sampling problem may lead to an underestimation of reaction rates.

In this study, the sampling time and number of trajectory frames are adjusted for the respective modelled reactions depending on the convergence of rates (see Supplementary Methods). The simulation setup is automated within KIMMDY and relies on user supplied simulation parameters. Simulation details for the molecular systems used in this study are described in the Supplementary Methods. KIMMDY is designed as a framework to be extended to diverse reactions. To this end, a plugin architecture providing a stable interface is available.

### 4.3 Choose reaction

From the collected event list, a reaction with a probability directly proportional to its rate is chosen. KIMMDY then associates with this event a time update from all predicted reactions. We implemented different kMC algorithms in a modular fashion in KIMMDY. In this work, only the rf-kMC algorithm with adaptive event list generation is used.

### 4.4 Effect reaction

To effect the chosen reaction, the corresponding reaction recipe is applied to change the molecular system topology, parameters and coordinates to the product state. Recipes define reactions through elementary recipe steps. Bind and break reference the two involved atoms and modify the MD bond definitions. Angles, dihedrals and pairs are modified accordingly. Changes of force field parameters are either handled by supplying a force field that has parameters for all reaction products or by re-parametrizing the bonded parameters with the generally applicable machine-learned Grappa force field[19] combined with heuristics for the nonbonded parameters (see Supplementary Methods). To generate the product coordinates, Place moves an atom in a certain snapshot to a new position and, as an alternative, Relax starts a MD simulation with the slow-growth feature of GROMACS to interpolate smoothly between reactant and product parameters. Finally, an equilibration MD simulation is performed to relax the new topology and reach an equilibrium ensemble as input for the next reaction sampling step. The equilibration timescale should be sufficient to ensure sampling for the next reaction from the product state Boltzmann distribution. Thus, KIMMDY models state transitions as a Markov process.

### 4.5 Efficiency of KIMMDY

With KIMMDY, simulations of consecutive and competing reactions in large biopolymers are possible for the first time. Taking the collagen fibril system with 2.6 million atoms as an example, tens of thousands of reaction rates are predicted per kMC step using physics-based or empirical models. For the machine-learned HAT model, about 100 reactions are evaluated every kMC step. While the necessary sampling and evaluation time depends on the rate prediction model, we observe that most time is spent on MD sampling for the aforementioned applications, and that the overhead introduced from the KIMMDY framework is negligible. Overall, one kMC step for the collagen fibril HAT application takes 10 h on a consumer GPU (Fig. A6a). Comparing this duration with calculating a single reaction barrier by DFT as done for the HAT GNN training, our emulator is roughly 100 times faster and at the same time accounts for conformational changes of the molecular system. Other systems see even greater speed-ups. With the hydrolysis reaction rates based on a heuristic, querying for possible reactions is done in seconds, rather than days.

### 4.6 Data availability

The datasets generated and/or analysed during the current study are available on GitHub at https://github.com/graeter-group/kimmdy-examples. Code that generates the simulations, results and figures is available at the same URL.

### 4.7 Code availability

KIMMDY is publicly available at https://github.com/graeter-group/kimmdy. The various KIMMDY plug-ins, *e.g.* for homolysis, hydrolysis, HAT and DNA crosslinking reactions, can be found under the kimmdy tag: https://github.com/topics/kimmdy.

## Acknowledgments

This work was supported by the Klaus Tschira Foundation (to F.G and K.R). This project has received funding from the European Research Council (ERC) under the European Union’s Horizon 2020 research and innovation programme (grant agreement No. 101002812) (to E.H., J.B. and F.G.). D.S and F.G gratefully acknowledge the support of the Klaus Tschira Foundation (SIMPLAIX project 8). D.S and F.G. gratefully acknowledge the provision of computing resources by the state of Baden–Württemberg through bwHPC and the German Research Foundation (DFG) through Grant Nos. INST 35/1597-1 (Helix cluster). We acknowledge funding through the Deutsche Forschungsgemeinschaft (DFG, German Research Foundation) under Germany’s Excellence Strategy – 2082/1 – 390761711. The authors gratefully acknowledge support by the German Research Foundation (DFG) grant GRK 2450. This research was conducted within the Max Planck School Matter to Life supported by the German Federal Ministry of Education and Research (BMBF) in collaboration with the Max Planck Society. The authors would like to thank Alexander I. Jordan and Alice Allen for helpful discussions and Alexander I. Jordan for suggestions regarding the statistical background of the methodology.

## 5 Author contributions

E.H.: Conceptualization, data curation, formal analysis, investigation, methodology, software, project administration, validation, visualization, writing(original draft, review & editing). J.B.: Conceptualization, data curation, formal analysis, investigation, methodology, software, project administration, validation, visualization, writing(original draft, review & editing). K.R.: Conceptualization, data curation, formal analysis, methodology, software, writing(review). E.U.: Formal analysis, investigation, methodology, visualization, writing (original draft, review & editing) B.N.S.: Formal analysis, investigation, methodology, software, visualization, writing (original draft, review & editing). D.K.: Formal analysis, investigation, software, visualization, writing (review & editing). D.S.: Data curation, formal analysis, investigation, methodology, visualization, writing (original draft, review & editing) C.A-S.: Conceptualization, Formal analysis, methodology, writing (review & editing) F.G.: Conceptualization, methodology, project administration, resources, supervision, writing (original draft, review & editing)

## 7 Materials & Correspondence

Correspondence and requests for materials should be addressed to F.G.

## Appendix A Extended Data

**Fig. A1.**
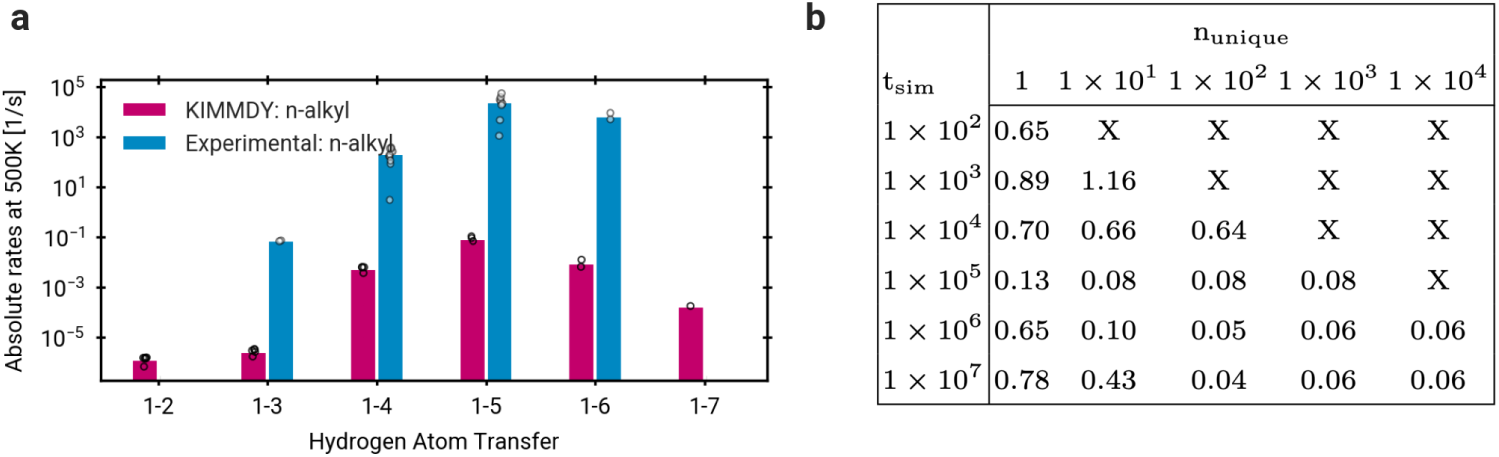
**a**, Absolute rates for all HAT reactions in all n-alkyl radicals (100 ns). Circles on the KIMMDY bar indicate mean of all simulations of a single n-alkyl radical type, e.g. 1-propyl. Circles on the experimental bar represent an extrapolated result for 500 K. Bars represent the mean over all circles.**b**, Mean Brier divergence of four runs for different combinations of number of MD frames in fs (rows) and number of barrier predictions per reaction *n unique*(columns) with the true probability distribution taken as the mean experimental rates at 500K for each shift. Note that the 1-2 and 1-7 HAT reactions have been excluded from the analysis, since there are no experimental values for comparison.

**Fig. A2.**
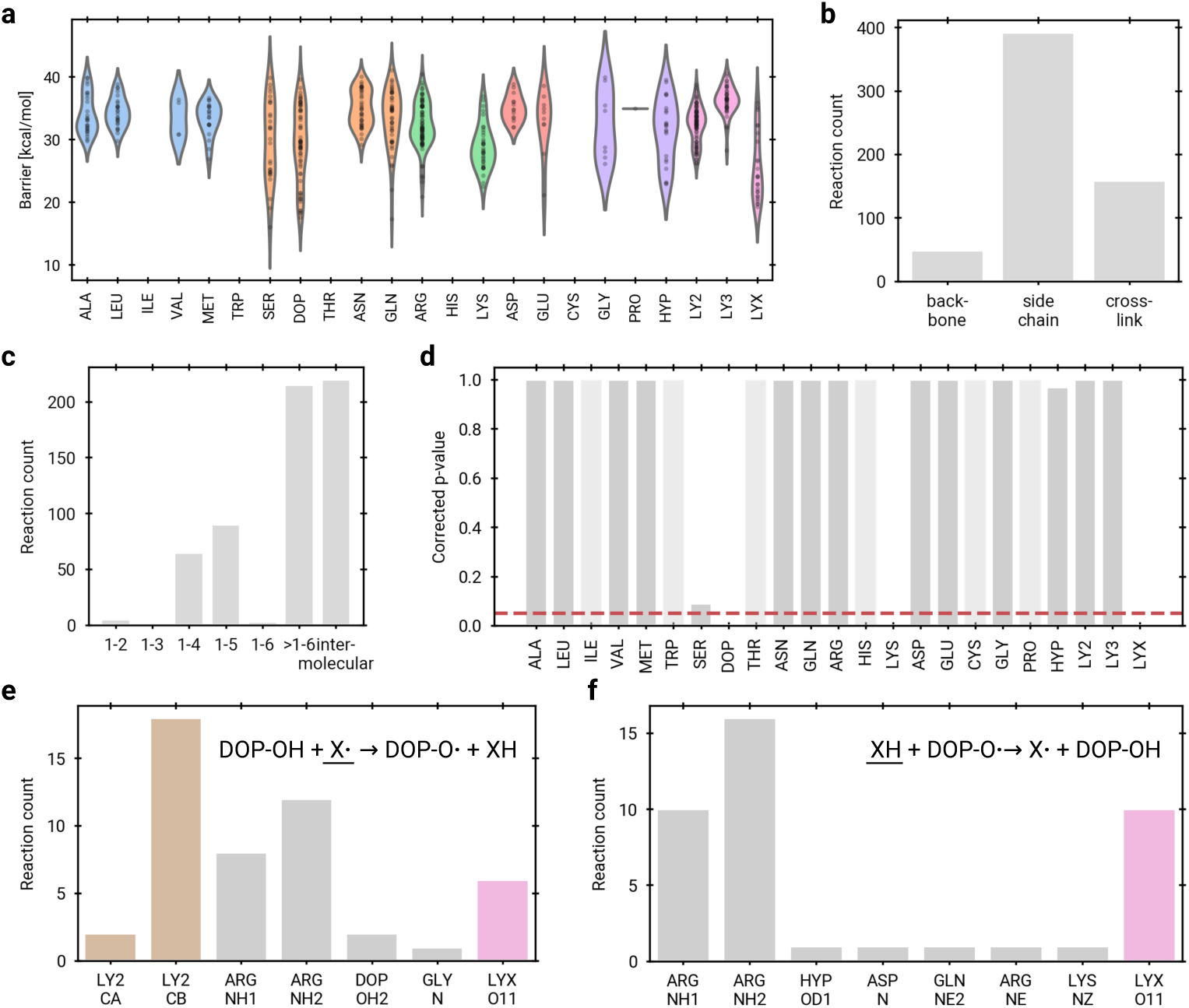
HAT occurs largely intermolecularly through side chains and a wide range of aminoacids. **a**, Barriers of HATs (n=586) by H-donor amino acids coloured by physicochemical properties. The PYD crosslink is coloured in pink and split into three amino acids LY2 LY3 and LYX with LYX containing the aromatic ring. **b**, Location of HAT H-donor classified by backbone, side chain and crosslink. **c**, Location of H-donors relative to H-acceptors. Atom numbering starts with the H-acceptors and is increased by 1 for every bond between donor and acceptor. **d**, Multiple-hypothesis testing of H-donors by amino acid with Welch’s t-test using the Benjamini and Hochberg approach[42] for controlling the false discovery rate. Amino acids with too few samples for testing are assigned a corrected p-value of 1 and shown in light grey. The red dashed line indicates the significance level of the corrected p-value at 0.05. **e**, Reaction count by identity of H-acceptors (underlined moiety in the inset scheme) for transfer with the DOPA hydroxy group as H-donor. Atoms names are following the Amber protein force field naming scheme. Atoms at the PYD short arm homolysis site are coloured in brown, atoms belonging to the aromatic ring of PYD in pink and all other atoms in grey. **f**, Reaction count by identity of H-donors (underlined moiety in the inset scheme) for transfer with the DOPA hydroxy group as H-acceptor.

**Fig. A3.**
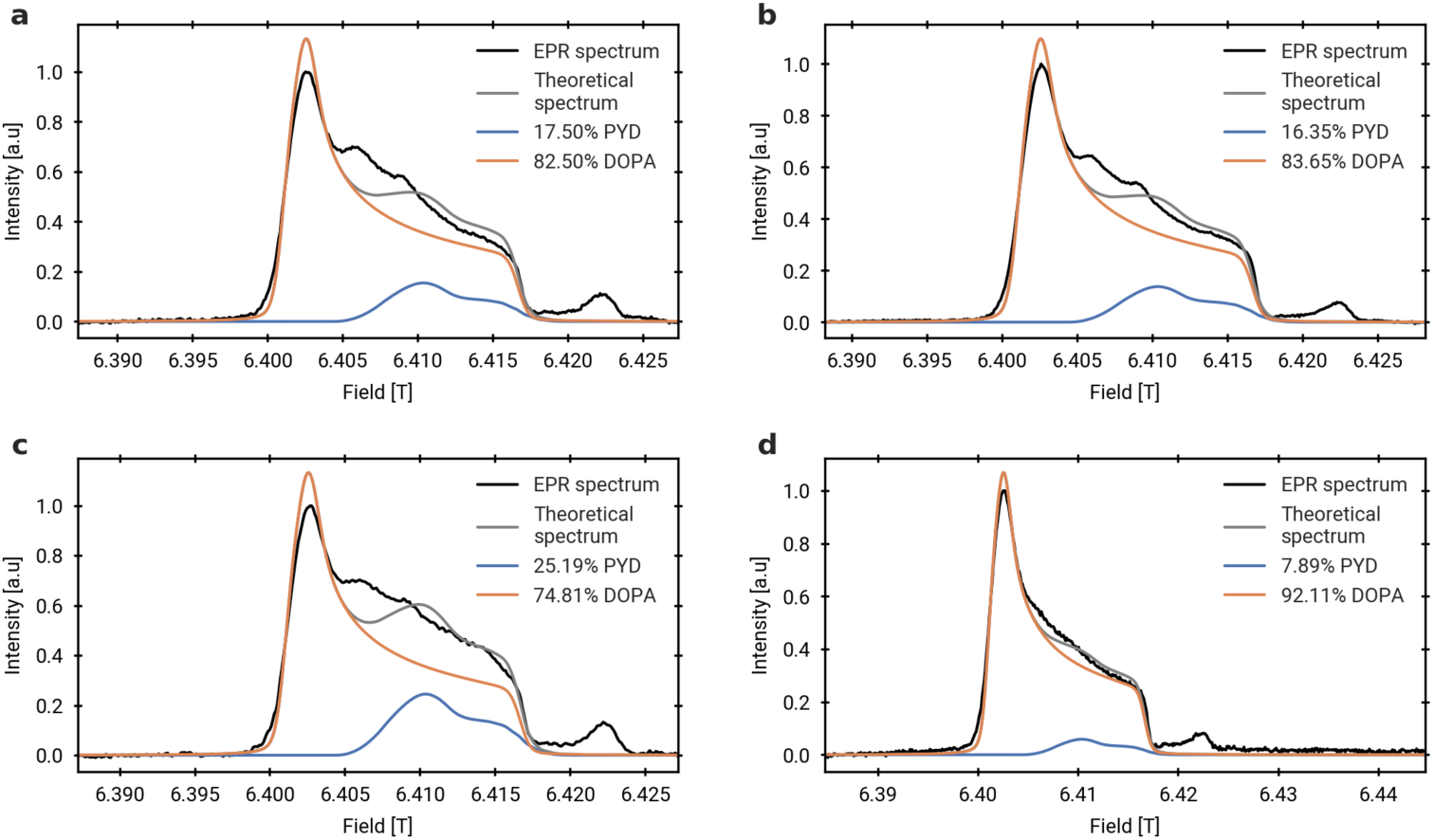
Collagen EPR spectra from different experiments. Panels a–c show experimental G-band EPR spectra of stretched collagen (black) reproduced from Kurth et al. [23], while panel (d) is taken from Zapp et al. [22]. Spectra from ab initio calculations are shown in comparison (in color), with the legend defining the relative contributions of DOPA and PYD used in the theoretical reconstructions.

**Fig. A4.**
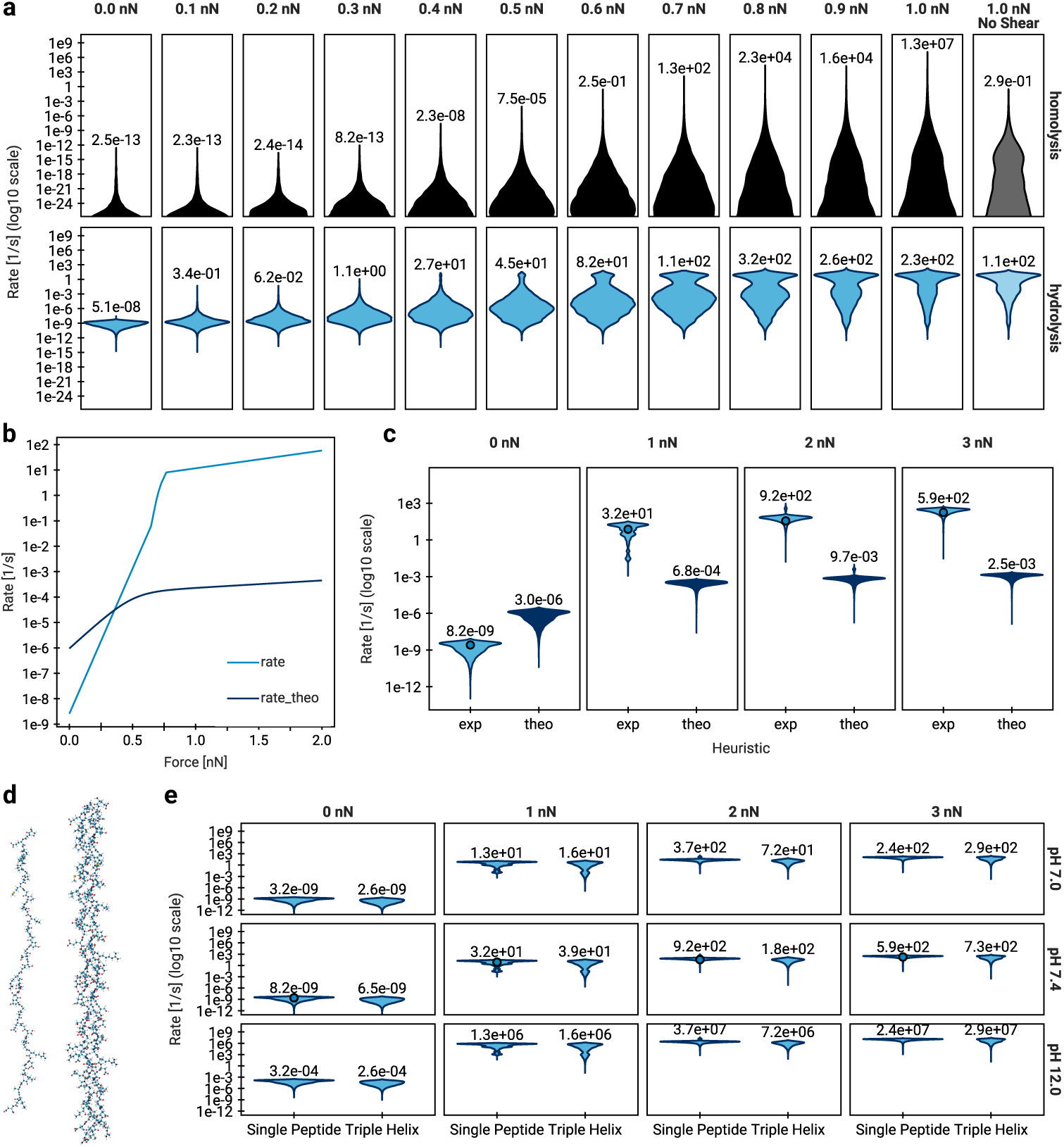
Analysis of hydrolysis reaction rates for additional systems and rate heuristics. **a**, Reaction rates of the homolysis and hydrolysis reactions in a collagen fibril pulled at increasing forces with a shear setup (see Supplementary Methods) compared to “No Shear” for the 1 nN. Uneven pulling, as would be found under physiological conditions, quickly ramps up the highest rates of each reaction due to force concentration in the fibril, while the overall bulk of rates is similar between the setups. But due to the nature of the random sampling and the vastly higher order of magnitude, the highest rate will dominate the result. Since the force dependence of the rates for hydrolysis (b) levels off sooner than for homolysis, homolysis is expected to prevail over hydrolysis at higher forces, especially in the shear setup. **b**, Force-response curve of the reaction rate of hydrolysis (“rate”) based on single molecule force spectroscopy experiments by [26] compared to the theoretical rate when calculated from reaction barriers from QM calculations by [26] (“rate theo”). **c**, Sampling of reaction rates of hydrolysis from simulations of the single peptide under increasing forces using the two aforementioned heuristics, theoretical from QM calculations and experimental from single molecule force spectroscopy. **d**, Molecular render of the single peptide and triple helix systems. **e**, The reaction rates of hydrolysis of the single peptide and triple helix systems under increasing forces for various pH values.

**Fig. A5.**
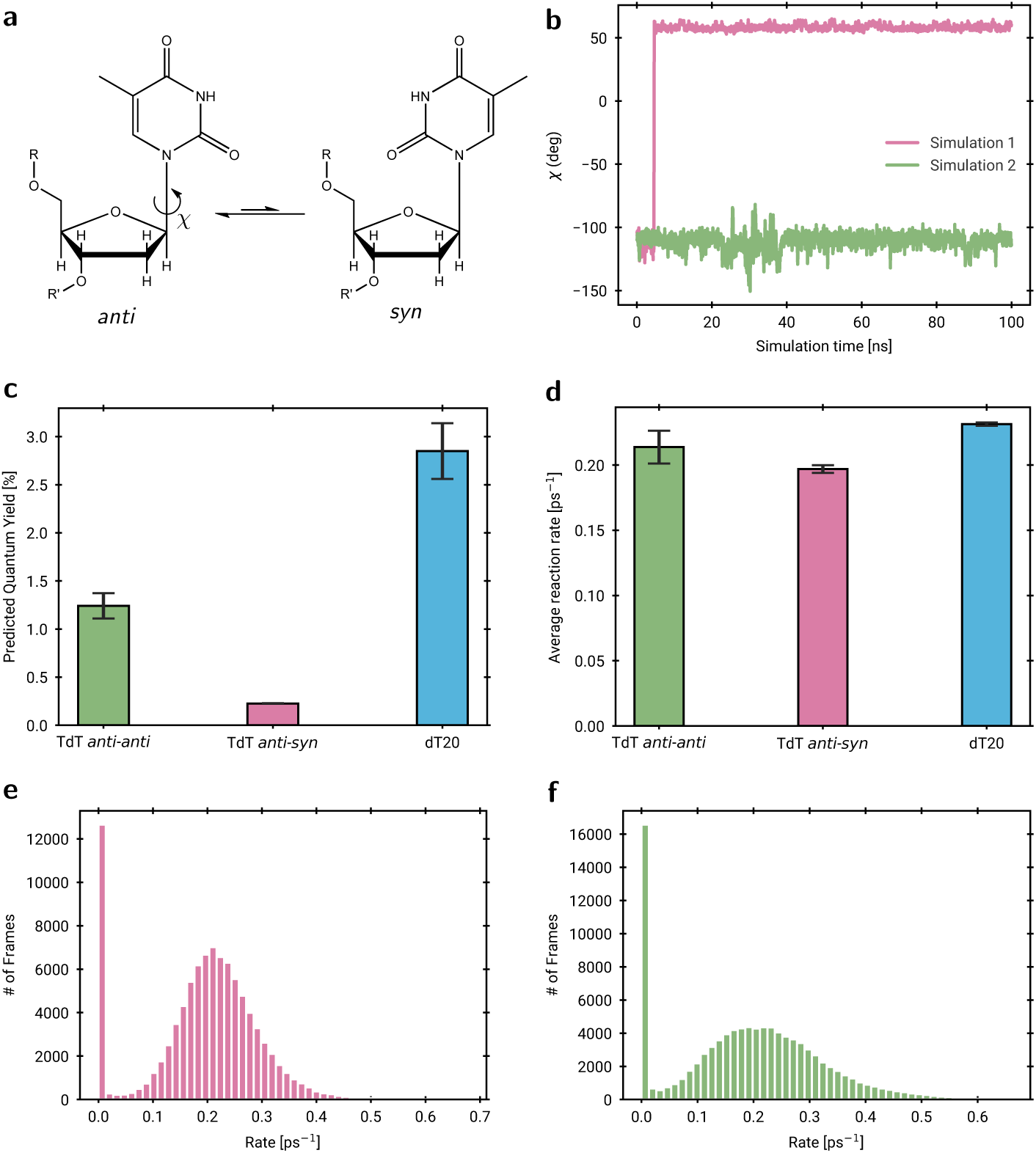
Conformational analysis in TdT systems. **a**, *syn-anti* equilibrium in thymine bases. **b**, Change of dihedral *χ* over the course of two simulations. Shown as running average of 100 frames for one of two thymines in the TdT system. Simulation 1 shows an *anti-syn* conformation, simulation 2 shows an *anti-anti* conformation. **c**, Quantum yields for benchmark systems (*n* = 4 for TdT *anti-anti*, *n* = 2 for TdT *anti-anti* and *n* = 3 for dT_20_ as separate 100 ns MD simulations). **d**, Average reaction rates for benchmark systems (equal number of samples as in c). **e**, Histogram of rates determined from one 100 ns TdT *anti-syn* simulation. **f**, Histogram of rates determined from one 100 ns TdT *anti-anti* simulation.

**Fig. A6.**
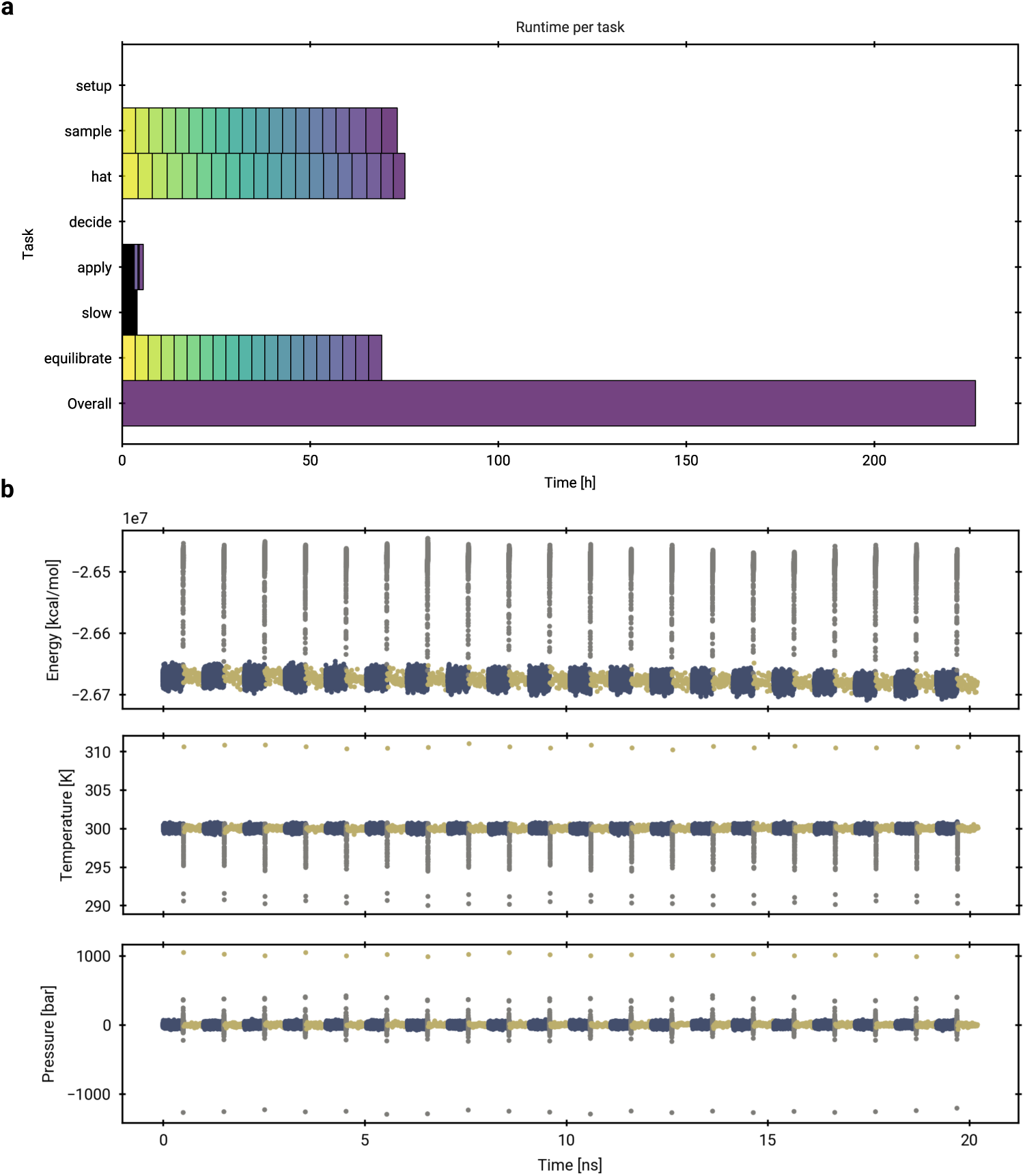
**a**, Elapsed real time of a collagen fibril KIMMDY run with 20 consecutive HAT reactions. Tasks are coloured using the viridis colormap by their call order. The tasks ‘sample’, ‘equilibrate’ and ‘slow’ are MD simulations, ‘hat’ is the HAT rate prediction, ‘decide’ is the choose reaction step and ‘apply’ the effect reaction step. **b**, Potential energy, temperature and pressure in the MD simulations of the same KIMMDY run as above. Sampling MD simulations are shown in blue, slow-growth parameter interpolation MD simulations in grey and equilibration MD simulations in pale yellow. Continuing the MD simulations before and after the slow-growth simulation creates artifacts from changing the type of simulation but the reference temperature and pressure are always reached again. Note that simulations are not continuous over reactions because the time to the next reaction passes according to the kMC algorithm.

