## Supplementary Information for "KIMMDY: a biomolecular reaction emulator"

### 1 Supplementary Methods

#### 1.1 KIMMDY Implementation Details

##### 1.1.1 Configuration, input/output

KIMMDY is controlled via a command-line interface (CLI) and a `yaml` formatted configuration file. In there, every task KIMMDY performs has a corresponding configuration section. The sequence of tasks KIMMDY executes is also defined in this configuration, making KIMMDY highly flexible. The configuration file contains input files such as the topology and initial coordinates, high performance computing (HPC) options, logging and working directory options, and general settings for KIMMDY.

For a complete documentation of all options, please read our extensive online documentation available at https://graeter-group.github.io/kimmdy/, which contains an automatically updated list of options at https://graeter-group.github.io/kimmdy/guide/references/input.html.

When the `yaml` file, typically named `kimmdy.yml`, is opened in a code editor supporting the language server protocol, such as the popular Visual Studio Code or Neovim, the user can get features such as auto completion for keywords and options as well warnings for missing or unknown keys. This reduces the likelihood of disappointing application crashes, because the validity of the input can be checked before KIMMDY is even started. Further consistency checks are front loaded at the start of KIMMDY such that it stops earlier rather than later in a run.

Additional provided tools include `kimmdy-analysis` for post-simulation analysis, and `kimmdy-modify-top` for modifying or parametrizing a system outside of a KIMMDY run.

##### 1.1.2 Ensemble Generation

In theory, KIMMDY can work with any ensemble generation method that results in atomistic trajectories. This includes *ab initio* MD, MLP simulations, classical MD or ensemble generation methods. In practice, the KIMMDY architecture is oriented towards GROMACS [1, 2] simulations, limiting the immediate uses to unbiased MD, biased MD, and in the near-future MLP simulations.

KIMMDY interfaces with GROMACS via its command-line interface, which makes KIMMDY robust to GROMACS version changes. In the KIMMDY config, the GROMACS binary name and flags for commands such as `gmx grompp` and `gmx mdrun` are exposed as options. The `.mdp` files for requested MD tasks and all files defining the molecular structure are taken as user input. It should be noted that albeit MD simulations are conducted, no continuous reactive trajectories are obtained. Rather, separate trajectories are generated for every state and the slow growth simulations (see below) in between. The concatenated trajectories that KIMMDY analysis tools can create from all states are only for visualization and inspection purposes.

##### 1.1.3 Reaction and Recipes

KIMMDY manages reactions via a plugin system, providing users with an easy method to expand KIMMDY's existing functionalities. A plugin generates reaction rates alongside reaction recipes based on the latest conformational ensemble. How the reaction rates are determined is up to the plugin. They can be calculated based on simple or complex physical models, predicted by a machine-learned model or be determined based on experimental heuristics. Plugins consume simulation outputs in various ways, but popular choices include using MDAnalysis [3, 4] to read trajectories or PLUMED [5] to monitor distances. The reaction recipes describe all modifications to the topology and structure required to perform the reaction corresponding to the calculated rate. A recipe is composed of recipe steps, which can break a bond, create a bond, place an atom at some coordinates, modify the topology, or request a specific MD simulation. Additionally, plugins can defer returning the actual steps of the recipe until a reaction has been chosen by KIMMDY to reduce computational cost. For example, the hydrolysis plugin returns rates for all peptide bonds in the system, but only chooses a specific water molecule to execute the attack once a bond has been chosen as the target. Reaction rates can be valid for the whole period of the sampling simulation or vary over time for each possible reaction, which is also reported back to KIMMDY.

##### 1.1.4 kMC algorithms

KIMMDY collects all reaction rates from the plugins and uses a user-selected choice of kinetic Monte Carlo (kMC) algorithm to sample a reaction from the distribution. For a theoretical background of kMC, please refer to Sec. 2.1.3. Choices for kMC algorithms currently include rejection free kMC ("rfkmc"), "extrande" [6], a modified version of extrande ("extrande_mod"), rejection free kMC with multiple choices at the same time ("multi_rfkmc"), as well as a debugging option that always chooses the first (or last) reaction. Individual reactions can declare their preferred algorithm in addition to the global choice of kMC algorithm. In this study, all systems use rejection free kMC.

##### 1.1.5 Topology module

From the initially user-supplied topology, an internal Topology object is build. The force field as well as topology includes are explicitly resolved, meaning every interaction has individual parameters. This enables the use of Grappa for parametrizing on-the-fly, but is still compatible with established force fields, like Amber [7] or Charmm [8]. The Topology supports breaking bonds, creating bonds, and deleting hydrogen atoms. For all of these actions, the interaction terms, like bonds, angles, dihedrals, pair exclusions etc., are updated accordingly. Furthermore, it identifies, and keeps track of possible radicals and whether it needs parametrization due to some modifications. Creating bonds, deleting bonds, and moving atoms does not change the index of the atoms stored in the topology. This ensures compatibility with standard analysis techniques, but residues are not necessarily continuous in the resulting topology afterwards. If continuous residues are needed, the Topology provides a function to re-index itself. After an atom is deleted, the topology is re-indexed by default. To finally use the Topology in an MD simulation, a new `top` file is generated.

##### 1.1.6 Parametrization interface

After each reaction, the product state needs to be parametrized. In KIMMDY, two options are available: basic parametrization and Grappa [9] parametrization (see Sec. 1.3.

Usage of the basic parametriser assumes that the product molecule parameters are contained in the force field included in the topology file, not modifying the topology beyond the instructions in the recipe.

Grappa parametrizes around the atoms involved in the reaction. The whole structure is parametrized with Grappa either before a KIMMDY run or at the start, producing consistent parameters for any reaction. A basic distinction defined in the KIMMDY config is between reactive and non-reactive molecules. Typically, solute atoms are defined as reactive and are consequently parametrized with Grappa. Solvent atoms are by default defined as non-reactive and have classical force field parameters, but can also be included in the reactive topology.

##### 1.1.7 Slow-growth

For creating an initial structure of the reaction product state, minimizing and equilibrating the molecular systems is viable. However, this may take a long time for large systems, resulting in inefficient simulations. We use the slow-growth method implemented in the GROMACS free-energy module to interpolate between product and reactant state for continuous parameter and coordinate changes. Subsequently, a short equilibration is necessary to ensure the states are Markovian.

Currently, exact continuations of MD simulations in GROMACS are only possible from checkpoint files, which do not allow for topology modifications. We use coordinates and velocities to continue MD simulations, which leads to a short period where thermostat and barostat equilibrate again. Switching between "free-energy" on and off in GROMACS also currently has some artifacts, albeit non-breaking, which may be improved in the future.

KIMMDY supports two modes of slow-growth. In "morse-only" mode, breaking or forming bonds are modelled entirely with morse potentials, where the state of the broken bond is mimicked by a morse potential with parameters such that the shape matches the Van der Waals potential of the two now non-bonded atoms. In "full" slow-growth mode all parameters around the affected atoms are transitioned, where non-bonded parameters are modelled using pair potentials. Slow-growth of non-bonded parameters, especially for Van der Waals interactions, can quickly lead to high energies and forces due to the exponential nature of the potential. As such the best choice of KIMMDY options and mdp parameters to relax the structure between reactions and sampling depends on the system. An example trajectory of a reaction including slow-growth can be seen at https://youtu.be/tgS5uleN288 and is visualized in Fig. S1.

##### 1.1.8 Checkpoints/restart

A checkpoint system facilitates long KIMMDY simulations (Fig. ??). Conveniently, GROMACS simulations can be restarted from checkpoints and the end of a simulation is a state where the current system state is written explicitly in files. Thus,

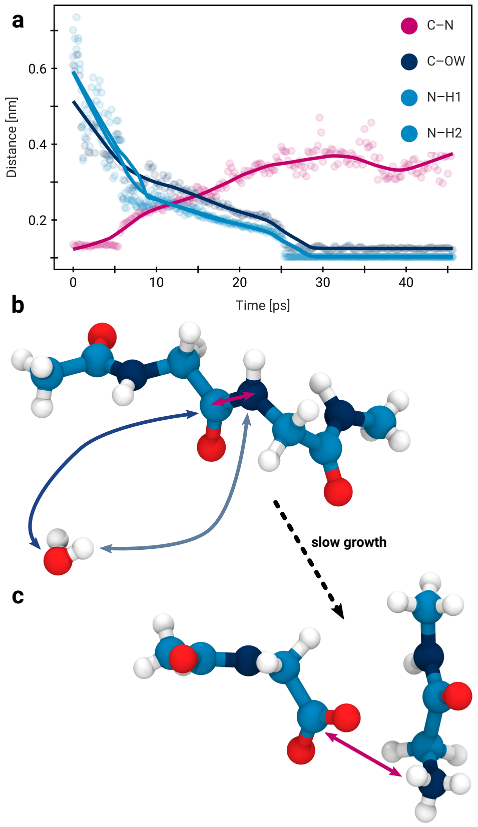

**Fig. S1 KIMMDY's slow growth applied to hydrolysis of a peptide bond**. **a**, A slow growth simulation visualized by the distances between the atoms involved in the reaction. **b**, Molecular renders of the peptide before and after (**c**) the reaction. Coloured double-arrows correspond to the distances in the plot.

we implemented KIMMDY restarting during and after GROMACS simulations. Files previously written by KIMMDY are parsed to faithfully reproduce the current state of the KIMMDY simulation. All tasks started after the last GROMACS task can not be used and are discarded. This is not typically an issue, because the simulations are the computationally expensive part, not the reaction rate queries.

##### 1.1.9 HPC interface

KIMMDY simulations are easily parallelizable and require extensive resources, typically available on High Performance Computing (HPC) systems. We added utilities to run KIMMDY on HPC systems using Slurm [10] and expose prefixes to the GROMACS commands necessary for running it on a system using a Message Passing Interface (MPI) such as OpenMPI. Restarting works for the HPC setting with automated resubmission of Slurm jobs that have a time limit.

##### 1.1.10 Code quality, Testing

Academic software developers often change rapidly due to the nature of short-term projects, such as Master's and doctoral degrees. Hence, writing robust and maintainable code was made a top priority in the development of KIMMDY. To minimize friction when working with multiple people on the same code base, we adhered to best practices of research software development and established clear conventions, outlined in our online documentation. Further, KIMMDY has infrastructure in place to automate running unit- and integration tests before a change is merged. New KIMMDY versions are automatically created based on the Conventional Commits specification to clearly identify new features and label backwards-incompatible changes.

##### 1.1.11 KIMMDY versions

The idea of running and restarting MD simulations after topology modification was first introduced in our group by [11] for homolytic bond breaks as a collection of scripts that were also called KIMMDY at the time. Subsequently, we set out to generalize it to arbitrary reactions, deepen the theory and make it available as a complete software package. We chose to keep the name KIMMDY and made the original KIMMDY available in the KIMMDY repository under the version 1.0.0. The current KIMMDY is now up to v8.0.0 at the time of submission.

#### 1.2 HAT GNN rate prediction

In this study, we use a previously developed graph neural network [12] to predict HAT reaction rates for an ensemble of structures generated from MD simulations. Reaction rates are predicted for individual conformations. The conformations are selected based on the distance between hydrogens and the radical, distances bigger than a defined cutoff get discarded. Further, conformations, for which steric clashes between the reacting hydrogen and its surrounding would occur during the reaction get discarded as well (as described in the original paper). For the prediction both the local reactant conformation and a putative product conformation with the hydrogen bound to the heavy atom radical are used as input. The GNN predicts barriers and corresponding rates are calculated with the Eyring equation.

The initial training was performed on more than 17 000 MD structures with corresponding DFT energy barriers with a MAE below 3 kcal/mol. A subset of these barriers were generated by optimizing the two heavy atoms and the involved hydrogen while keeping other atoms frozen. Then, a gradient-based transition state search was performed. The resulting rates correspond to instantaneous transition rates for a given structure. Thus, predicting HAT rates for an ensemble of structures yields only rates for reactions that could occur given the conformational constraints of the surroundings instead of the minimum energy path for unconstrained fragments. Nonetheless, since the environment is constrained, this could lead to overestimated barriers resulting in an underestimation of reaction rates.

The previously published HAT barrier model was trained on structures sampled from MD simulations based on the amber99SB-ILDN [13] force field. A central part of KIMMDY is the reparametrisation of systems on-the-fly using Grappa. However, this reparametrisation leads to different input structures for the HAT model, as Grappa generates proper radical parameters, which are absent in the traditional force fields. To predict HAT barriers on structures parametrized with Grappa, a HAT barrier models trained on such structures is needed. Hence, we use a new version of the HAT barrier model suited for Grappa structures (Fig. S2). It was trained in a transfer-learning scheme, analogous to the transfer learning described in the previous version.

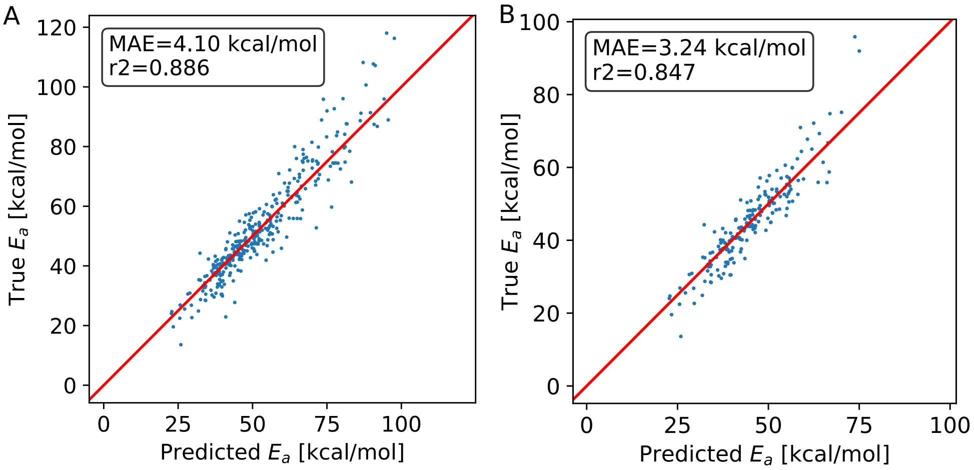

**Fig. S2 Performance of the HAT model trained on Grappa structures. a**, 360 structures $< 3\AA$ and **b**, Structures $< 2\AA$ translation distance of the reacting H Atom.

HAT rates correlate strongly with the distance a hydrogen moves between reactant and product state, called translation distance (Fig. S7a). Due to the exponential nature of the barrier-rate relation, a limited amount of rates contributes a large fraction of the sum of all rates. Thus, the structures with the lowest translation distances per reaction are expected to suffice to approximate the mean rate of a reaction. The exact number of evaluated structures per reaction is a hyperparameter and should be screened for every new application. This HAT rate prediction approach has been newly implemented for this work to facilitate KIMMDY simulations of large biomacromolecular systems. Further runtime improvements involve optimized trajectory processing using MDAnalysis [14, 15].

#### 1.3 Grappa

Grappa [9] is trained on molecule-wise relative QM energy and forces to predict sets of classical force field bonded parameters. As input features, it uses only the atomic numbers, partial charges, number of neighbours and ring membership, making it suitable for parameter predictions of diverse molecules at state-of-the-art accuracy. Parameters do not depend on the molecule conformation. Grappa force fields can be used in established MD engines, simulating several orders of magnitude faster than MLIPs such as Allegro and MACE [16, 17].

The chemical space covered by Grappa includes small molecules, peptides and RNA [18, 19]. The parameters for peptides are validated against a range of NMR J-couplings, outperforming current protein force fields. Apart from the canonical amino acids, Grappa also predicts parameters for hydroxyproline and DOPA, which are abundant in collagen, as well as peptide radicals.

Reaction products in KIMMDY simulations are efficiently parametrised with Grappa. At the beginning of a simulation, molecular systems are parametrised in their entirety. After reactions, only a region around the atoms involved in the reaction is reparametrised. In this study, the reparametrised region includes atoms up to 7 bonds away from the reacting atoms. The field-of-view is twice as far, *i.e.* 14 bonds so that all reparametrised atoms receive information about their neighbouring atoms. Atoms further away from the reacting atoms are not influenced by the reaction because current Grappa models pass messages up to four bonds away. Note that classical biomolecular force fields parametrise amino acids independent of their neighbouring amino acids, hence their field-of-view is smaller than Grappa's.

Grappa bonded parameters are combined with Amber99 nonbonded parameters for proteins and GAFF2 nonbonded parameters for small molecules [20]. For reactions, charges are modified to keep fragments at integer charge. This is done by moving charge on atoms involved in reactions and keeping other charges unchanged.

| Dataset | Test Mols | Confs | RMSE | Grappa | Grappa-radical | Mean predictor |
|---|---|---|---|---|---|---|
| SPICE-Pubchem | 1411 | 60853 | *Energy* | **2.3** | 2.4 | 18.4 |
| | | | *Force* | **6.1** | 6.4 | 23.4 |
| SPICE-DES-Monomers | 39 | 2032 | *Energy* | **1.3** | 1.4 | 8.2 |
| | | | *Force* | **5.2** | 5.5 | 21.3 |
| SPICE-Dipeptide | 67 | 2592 | *Energy* | **2.3** | 2.5 | 18.7 |
| | | | *Force* | **5.4** | 5.7 | 21.6 |
| RNA-Diverse | 6 | 357 | *Energy* | **3.3** | 3.5 | 5.4 |
| | | | *Force* | **3.7** | 4.4 | 17.1 |
| RNA-Trinucleotide | 64 | 35811 | *Energy* | **3.5** | 3.7 | 5.3 |
| | | | *Force* | **3.6** | 4.3 | 17.7 |
| Peptide-Radical-Opt | 17 | 818 | *Energy* | | **3.7** | 6.9 |
| | | | *Force* | | **9.1** | 7.7 |
| Peptide-Radical-Scan | 1 | 1235 | *Energy* | | **2.9** | 5.0 |
| | | | *Force* | | **4.9** | 3.9 |
| Peptide-Radical-MD | 16 | 1598 | *Energy* | | **2.4** | 5.3 |
| | | | *Force* | | **8.7** | 19.7 |

**Table S1** Accuracy of Grappa and Grappa-radical on MD sampled datasets used for training Grappa-1.4.0 and on peptide radical datasets. States in all datasets were MD sampled at temperatures of 300 K or 500 K. RMSEs of zero-centred energies are in kcal/mol, component-wise RMSEs of forces in kcal/mol/Å. SPICE-Pubchem is a small-molecule dataset, SPICE-DES-Monomers and SPICE-Dipeptide peptide datasets and RNA-Diverse and RNA-Trinucleotide are RNA datasets. The $\omega$B97M-D3(BJ) functional and def2-TZVPPD basis set were used for non-radical data, BMK/6-311+G(2df,p) for radical data.

In this study, the model Grappa-1.4.1-radical is used for parametrisation. Compared to the published model 1.4.0, it has been trained on the original data set plus peptide radical data to further improve the accuracy on radicals created by homolysis or hydrogen abstraction (See Table S1). All hyperparameter settings are the same as in the original Grappa publication.

#### 1.4 Bond Dissociation Energies

The bond dissociation energy (BDE) quantifies the enthalpy required to homolytically cleave a bond [21]. The lower the BDE, the higher the likelihood of bond rupture due to thermal fluctuations. In the case of abstraction reactions on a molecule X, the BDE is defined as

$$BDE(\mathrm{X}) = H(\mathrm{H}^\bullet) + H(\mathrm{X}^\bullet) - H(\mathrm{X\text{-}H}) \ , \tag{1}$$

where $H$ is the enthalpy, and the dot represents the presence of a radical. While straightforward, this direct approach may accumulate systematic errors, particularly those associated with open-shell species [22]. A clever way to overcome this problem is to compute the BDEs using the isodesmic method ($\mathrm{BDE}_{iso}$) [23], which yields more reliable values than the ones given by the equation 1. The $\mathrm{BDE}_{iso}$ is obtained by using a known BDE experimental value, $\mathrm{BDE}_{exp}$, of a reference molecule, Y, and replacing the enthalpy of the hydrogen radical in equation 1, which yields

$$BDE_{iso}(\mathrm{X}) = [H(\mathrm{X}^\bullet) - H(\mathrm{Y}^\bullet)] + [H(\mathrm{Y\text{-}H}) - H(\mathrm{X\text{-}H})] - BDE_{exp}(\mathrm{Y}) \ . \tag{2}$$

Note that the first and second brackets of the equation 2 contain substractions of exclusively open-shell and closed-shell species, respectively. In this way, systematic errors related with spin contamination are reduced instead of being accumulated. The same logic is followed for the cancellation of other systematic errors in isodesmic methods, which then works better when the molecule X and Y are similar in terms of the kind and number of bonds [22].

For the computation of the BDEs of DOPA and PYD shown in this paper (Fig. 2), we use phenol as the reference molecule, which has an experimental BDE value of 86.711 kcal/mol [24]. The structure of the PYD molecule is shown in Fig. S3. All the calculations of enthalpies were obtained using the Orca 5.0 software [25, 26], employing the M06-2X/def2-TZVP level of theory, which is parametrised to perform well in thermochemistry and kinetics applications [27]. We compare our computed values with the BDEs reported by Treyde *et al.* [28], who also used the isodesmic method for a large set of amino acids.

For simplicity, we use the notation "BDE" instead of "$\mathrm{BDE}_{iso}$" to refer to the bond dissociation energies computed using the isodesmic method in the other sections of this paper.

#### 1.5 Electron Paramagnetic Resonance spectrum

Electron Paramagnetic Resonance (EPR) spectroscopy leverages the splitting of the electronic levels of energy when a magnetic field is applied. That separation of the

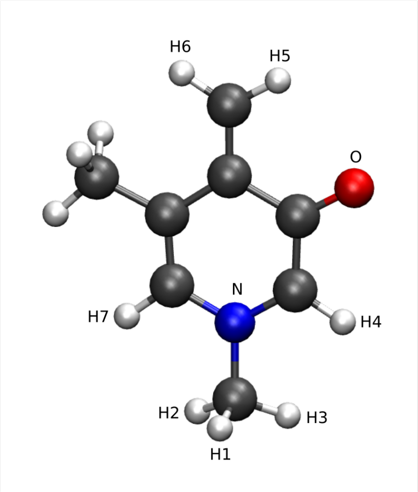

**Fig. S3** Render of the central part of the PYD crosslink used for the calculations of the electronic structure. The labels correspond to the atoms considered to compute the hyperfine coupling.

levels of energy allows observing electronic transitions by light absorption in systems with unpaired electrons (Zeemann effect). The magnitude of the separation of the levels of energy relates with the intensity of the magnetic field, **B**, is described by the spin Hamiltonian, defined as [29]

$$\hat{H}_{spin} = \mu \mathbf{B}\mathbf{g}\hat{\mathbf{S}} + \sum_i [\hat{\mathbf{S}}\mathbf{A}^{(i)}\hat{\mathbf{I}}^{(i)}] \ , \quad (3)$$

where $\hat{\mathbf{S}}$ and $\hat{\mathbf{I}}$ are the electronic and nuclei spin, respectively, **g** is the g-tensor, **A** is the hyperfine coupling tensor, and $\mu$ is the electronic Bohr magneton. The absorption spectrum can be computed by diagonalizing the spin Hamiltonian [30]. **g** and **A** are molecular parameters, obtained from the electronic density.

For the PYD EPR spectrum presented in this paper (Fig. 2) , we compute the spin-Hamiltonian parameters with the Orca 5.0 package [25, 26], employing the GIAO gauge and the B3LYP/EPRII level of theory in order to reduce the dependency with the system of coordinates [31, 32] (table S2 and S3). We use the same parameters of DOPA used by Kurth *et al.* [33]. Once the EPR parameters are computed, we obtain the EPR spectrum using EasySpin [30]. The theoretical spectrum is obtained as a linear combination of both spectra,

$$S_{theoretical}(B_0) = c_1 \ S_{DOPA}(B_0) + c_2 \ S_{PYD}(B_0) \ , \quad (4)$$

where the coefficients $c_1$ and $c_2$ are optimized to fit the experimental data. Given that the intensity is arbitrary, we can normalize the coefficients in order to have an estimate of the percentage of DOPA and PYD. In the figure presented in the main text, the percentages were 82.5% for DOPA, and 17.5% for PYD. However, those percentages change in independent measurements of different tissues (as shown in the Fig. **??**), because the concentration of each radical differs from sample to sample.

Although in the theoretical computation of the EPR spectra the hyperfine interactions are often neglected, we note that the EPR spectrum of the PYD crosslink is highly affected by including this correction (Fig. S4 and Table. S3).

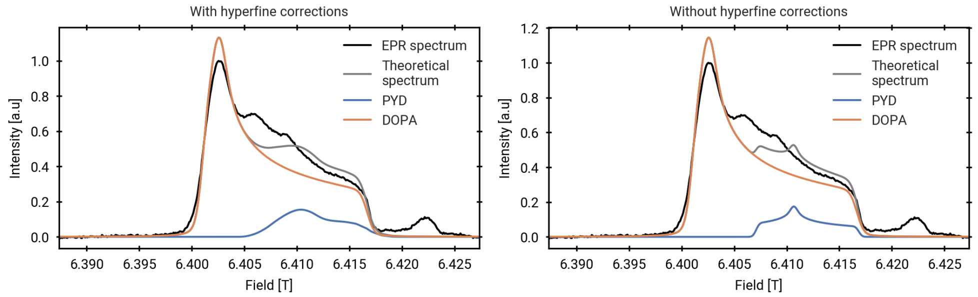

**Fig. S4** EPR spectra with (left) and without (right) hyperfine interactions included in the theoretical reconstruction.

| $\mathbf{g}_x$ | $\mathbf{g}_y$ | $\mathbf{g}_z$ |
|---|---|---|
| 2.0022 | 2.0066 | 2.0071 |

**Table S2** g-values of PYD

| | $\mathbf{A}_x$ | $\mathbf{A}_y$ | $\mathbf{A}_z$ | $\mathrm{A}_{iso}$ |
|---|---|---|---|---|
| N | -0.8717 | -1.2717 | 23.3633 | 7.0733 |
| O | 15.2727 | 15.7918 | -71.3462 | -13.4272 |
| H1 | 20.5010 | 20.7204 | 25.6272 | 22.2829 |
| H2 | 2.7320 | 3.0484 | 8.0320 | 4.6041 |
| H3 | 4.3020 | 4.5235 | 10.0596 | 6.2950 |
| H4 | -9.3574 | -26.3606 | -38.1397 | -24.6192 |
| H5 | -10.3160 | -25.8253 | -39.4415 | -25.1942 |
| H6 | -12.5424 | -27.1301 | -38.2859 | -25.9861 |
| H7 | 4.4109 | 5.1462 | 11.3476 | 6.9682 |

**Table S3** Diagonal hyperfine tensors for the selected niclei showed in Fig. S3. All the values are given in MHz.

#### 1.6 Rates in DNA dimerisation

For the TdT system, we determine whether a simulation samples the *anti-anti* or the *syn-anti* conformation by examining the rotation of the N-glycosidic bond characterised by the dihedral angle $\chi$ (see Fig. A5a). We find that some simulations switch to the *syn* conformer at one of the thymines early in the simulation and never return to the *anti* conformer (see Fig. A5b). To determine quantum yields for simulations that switched to the *syn-anti* conformation, we truncate the trajectory from the frame in which the switch happened. Quantum yields for the TdT and the $dT_{20}$ systems neatly match the experimental values, as is to be expected since the parameters of the model have been determined using those values (see Fig. A5c). Additionally, the *syn-anti* conformation shows a significantly lower quantum yield, as discussed in the main text. The average reaction rates of the three systems (TdT *anti-syn*, TdT *anti-anti*, and $dT_{20}$) show similar values (see Fig. A5d). Finding different quantum yields but comparable average reaction rates can be explained by examining the distributions of rates (see Fig. A5e,f), which show that the TdT *syn-anti* conformer has fewer very low rates but also cannot consistently reach rates above 0.5 $ps^{-1}$. TdT *anti-anti* reaches high rates above the threshold more often but also shows more frames with very low rates, indicating an increased flexibility in both directions regarding the reaction coordinate of thymine dimerisation.

KDE density plots for ds center, nicked, overhangs, and crossover systems are shown in Fig. S5a–d. The KDEs were computed from 10000 frames randomly selected across all MD simulations and normalized to their respective maxima.. We find that the increased flexibility of crossovers, as discussed in the main text, is also observed from overhangs, which seem to be less flexible than crossovers but still more flexible than ds center or nicked systems. Additionally, we find a decrease in average rate for overhangs and crossovers (see Fig. S5e). The systems for consecutive dimerisation reactions show no difference in average rates (see Fig. S5f).

#### 1.7 Simulation Details

##### 1.7.1 n-alkyl radicals

***Structure generation***

The n-alkyl molecule structures were generated and minimized using OpenBabel [34]. Antechamber24 [35, 36] was used to generate GAFF2 [7] parameters and were converted to GROMACS topologies using AcPype [37]. The radicals were generated via the KIMMDY modify_top tool and parametrized via Grappa.

***Simulation***

We simulated each alkyl radical using stochastic dynamicsfor 100 ns with 1 fs steps. Every 100 fs a conformation was added to the set of possible reactions for a total of $10^6$ conformations for further prediction. For each possible reaction pair we filtered for the 500 reactions (see section 1.2) with the smallest translation distance. For these reactions, the barriers were predicted using the ensemble of HAT GNN models. The rates were calculated using the Eyring equation as described in section 2.2. All simulations were repeated 20 times with different initial seeds.

We chose to exclude reactions of primary carbon to the other primary carbon from the analysis, since these cannot be measured experimentally. For comparing the selection probability, we only used simulations of 1-heptyl and 1-octyl radicals, since these are the only systems that include all considered reactions, i.e. 1-2 to 1-6 shifts in

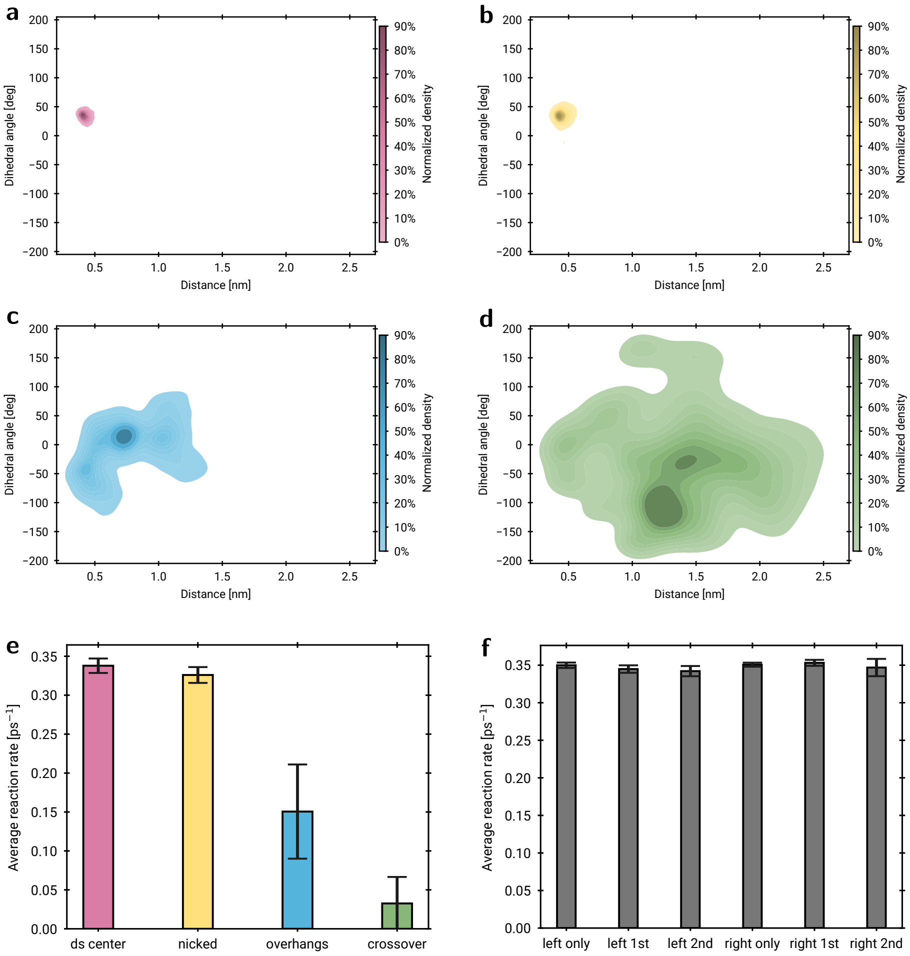

**Fig. S5 KDE densities and reaction rates for complex DNA systems. a-d**, Kernel density estimate (KDE) of distance and angle distributions of ds center (a), nicked (b), overhangs (c) and crossover (d) systems determined from 10000 snapshots randomly sampled from all MD simulations available for the respective system. The distances are the distances between reactive thymine double bonds and the angles are dihedrals within two thymine double bonds. **e**, Average reaction rates for common DNA origami motifs ($n = 3$ for ds center, nicked, and overhangs, and $n = 6$ for crossover from separate 100 ns MD simulations). **f**, Average reaction rates consecutive dimerisation systems (and respective control systems), in which the two reactive sites are five base pairs apart ($n = 3$ for left only, left 1st, right only, right 1st, $n = 4$ for right 2nd, and $n = 5$ for left 2nd from separate 100 ns MD simulations).

1-heptyl and additionally the 1-7 shift for 1-octyl. Overall, the isomerisation reaction classes in n-alkyl radicals should be considered to be comparable over n-alkyl radicals over different chain lengths, e.g. 1-5 in 1-hexyl and 1-heptyl should have similar barriers [38] and rates [39].

##### *Experimental rates*

In table S4 we summarize the experimentally derived alkyl HAT rates used in the main text. The experimentally obtained rate equations were extrapolated to 500K. This temperature was chosen to be closer to the typical range of experimental results. Note that due to this extrapolation outside their measured or indicated temperature range, the experimental rates will have an inherent uncertainty, in addition to the conventional experimental error.

| shift | n-alkyl radical | rate equation $[1/s]$ | rate at 500K $[1/s]$ | temperature range [K] | source |
|---|---|---|---|---|---|
| 1-3 | pentyl | $k = 4.10944717823092 \cdot 10^{-12} T^{6.837} e^{-\frac{9444}{T}}$ | $7.3 \times 10^{-2}$ | 700-1900 | [40] |
| 1-3 | pentyl | $k = 3.8904514499428 \cdot 10^{-12} T^{6.843} e^{-\frac{9451}{T}}$ | $7.1 \times 10^{-2}$ | 400-1900 | [41] |
| 1-4 | pentyl | $k = 488000000.0 T^{0.846} e^{-\frac{19.53}{RT}}$ | $2.7 \times 10^{2}$ | 350-1300 | [42] |
| 1-4 | hexyl | $k = 6.98 T^{3.2} e^{-\frac{8333}{T}}$ | $1.7 \times 10^{2}$ | 500-1900 | [43] |
| 1-4 | heptyl | $k = 117.489755493953 T^{2.85} e^{-\frac{8680}{T}}$ | $1.7 \times 10^{2}$ | 860-1050 | [44] |
| 1-4 | pentyl | $k = 330000000.0 e^{-\frac{15.1}{RT}}$ | $8.3 \times 10^{1}$ | 297.15-435.15 | [45] |
| 1-4 | hexyl | $log(k) = 11.0 - \frac{10.4347826086957}{RT}$ | 3.1 | 723-823 | [46] |
| 1-4 | pentyl | $k = 14000000.0 e^{-\frac{10800.0}{RT}}$ | $2.7 \times 10^{2}$ | 438.5-502.5 | [47] |
| 1-4 | pentyl | $log(k) = 11.08 - \frac{8.70169344333478}{RT}$ | $2.1 \times 10^{2}$ | 438-923 | [48] |
| 1-4 | pentyl | $k = 2432.0 T^{2.324} e^{-\frac{8183}{T}} + 911100.0 e^{-\frac{5303}{T}}$ | $3.8 \times 10^{2}$ | 300-1300 | [49] |
| 1-4 | octyl | $k = 5.12861383991365 T^{3.23} e^{-\frac{8479}{T}}$ | $1.2 \times 10^{2}$ | 700-1900 | [50] |
| 1-4 | pentyl | $k = 11.8338134809806 T^{3.03} e^{-\frac{7696}{T}}$ | $3.7 \times 10^{2}$ | 700-1900 | [40] |
| 1-4 | pentyl | $k = 11.4815362149688 T^{3.033} e^{-\frac{7706}{T}}$ | $3.6 \times 10^{2}$ | 400-1900 | [41] |
| 1-5 | hexyl | $log(k) = 9.41 - \frac{4869.5652173913}{RT}$ | $3.2 \times 10^{4}$ | 298.15-378.15 | [51] |
| 1-5 | hexyl | $ln(k) = 9.5 \log(10) - \frac{11.6}{RT}$ | $2.7 \times 10^{4}$ | 300-453 | [52] |
| 1-5 | hexyl | $k = 66500000.0 T^{0.823} e^{-\frac{12.45}{RT}}$ | $4.0 \times 10^{4}$ | 350-1300 | [42] |
| 1-5 | hexyl | $k = 183.0 T^{2.55} e^{-\frac{5516}{T}}$ | $2.3 \times 10^{4}$ | 500-1900 | [43] |
| 1-5 | heptyl | $k = 676.082975391982 T^{2.39} e^{-\frac{5237}{T}}$ | $5.4 \times 10^{4}$ | 860-1050 | [44] |
| 1-5 | hexyl | $log(k) = 10.5 - \frac{7.39130434782609}{RT}$ | $1.1 \times 10^{3}$ | 723-823 | [46] |
| 1-5 | hexyl | $k = 20000000.0 e^{-\frac{8300.0}{RT}}$ | $4.7 \times 10^{3}$ | 350-410 | [53] |
| 1-5 | octyl | $k = 22.9086765276777 T^{2.82} e^{-\frac{5413}{T}}$ | $1.9 \times 10^{4}$ | 700-1900 | [50] |
| 1-6 | heptyl | $k = 245.470891568503 T^{2.51} e^{-\frac{6292}{T}}$ | $5.0 \times 10^{3}$ | 860-1050 | [44] |
| 1-6 | octyl | $k = 2.95120922666639 T^{3.08} e^{-\frac{5544}{T}}$ | $9.3 \times 10^{3}$ | 700-1900 | [50] |

**Table S4** Experimental rates used for n-alkyl isomerisation in the main text. Here we use constant $R = 1.987 \times 10^{-3}$ kcal/mol/K.

##### *Quantitative comparison of reaction probabilities using the Brier divergence*

To quantify the discrepancy of the selection probabilities, we use the Brier divergence (BD) [54],

$$d_B(P, Q) = \sum_{i=1}^{k} (p_i - q_i)^2, \tag{5}$$

where $P$ and $Q$ are the set of predicted and true categorical probabilities $\{p_i\}$ and $\{q_i\}$, respectively and $k$ is the number of categories or shifts. The divergence is minimized when $P = Q$ with value 0 and maximized with value 2 when $p_i = 1$ and $q_j = 1$ for $i \neq j$. Consequently, overconfident predictions are penalized stronger than an ignorant uniform prediction over all categories.

In the case of the results of the main text, the BD for an ignorant uniform prediction is 0.48. The mean BD of the prediction distributions obtained using the 1-heptyl and 1-octyl is 0.04 and 0.02, respectively. The low scores confirm the strong agreement of the distributions seen by visual inspection.

***Simulation hyperparameters***

The two main parameters for the KIMMDY simulations with the HAT GNN are the total simulation run *t_sim* and the maximum number of barriers *n_unique* to be predicted per reaction (see section 1.2) during the MD phase. E.g. *n_unique* of 100, would mean that per reaction pair, the frames with the 100 smallest reaction distances would be used for barrier prediction. For the rest of the frames the barriers are assumed to be so high that the reaction rate should be zero.

The mean BD of different combinations of *t_sim* and *n_unique* can be seen in Fig. A1 b. Note that in this case, we assume sufficient decorrelation of barriers between time steps and therefore do not simulate for shorter times. Instead, we subsample from the longest run used in Fig. 2a, i.e. we take every 10th step to get an expected simulation time that is ten times shorter than the longest run; every 10th step of that shortened run to get a run ten times shorter again etc.

The convergence of the mean rate for each shift for a 10 ns run, sampled by taking every 10th frame from the original 100 ns runs as described above, is shown in Fig 2 d with different *n_unique* values on the horizontal axis. It can be seen that the rates have mostly converged between 100 and 1000 number of predicted reactions. The 1-4 reaction becomes more likely compared to the 1-6 reaction after using 1000 barrier predictions. We therefore used 500 as a compromise between accuracy and practical efficiency for a realistic use case. Notable exceptions are the 1-2 and 1-3 transitions, which have steadily rising reaction rates. This is probably because the distance between the radical centre and the hydrogen atom will be relatively similar and the barriers at almost all simulation steps are non-negligible. Nonetheless, the barriers are so high in these case, that they are non-competitive with respect to the other transitions.

***Ring strain energy***

HAT isomerisation reactions in n-alkyl radicals occur through ring structured transition states, and the barrier of an 1-$x$ HAT is often approximated using $E_{abs} + E_{RS}$, where $E_{RS}$ is the ring strain energy associated with the $x + 1$ ring-like transition state structure and $E_{abs}$ is the barrier for a HAT reaction in a bimolecular case [38]. The ring strain is largest for the 1-2 case and decreases until 1-5. For reactions of 1-6 and larger ring sizes, entropic effects are thought to play a stronger role leading to a decrease in reaction rates [39].

##### 1.7.2 Radical Migration in Collagen

Simulations of consecutive HAT reactions are conducted with an all-atom model of a *Rattus norvegicus* collagen fibril (PDB ID: 3HR2) comprised of 41 triple helices spanning one central overlap and one gap region for a total of roughly 320 000 protein atoms. ColBuilder [55] was used to generate a model with N- and C-terminal PYD crosslinks with a connectivity of 9.C-5.B-944.B and 1047.C-1047.A-98.B, respectively. Furthermore, all Phe and Tyr residues were randomly mutated to either DOPA deprotonated at the C$\epsilon$ (DO1) or C$\zeta$ (DO2) hydroxy group. Since no quantitative experimental data of the oxidation state of Phe and Tyr residues has been published, these mutations serve to treat all potential DOPA sites as equally strong radical scavengers.

The fibril was solvated, salt ions were added to neutralize the system and reach a concentration of 150 mM. After an energy minimization using the steepest descent method, a 10 ns NVT equilibration and then a 10 ns NpT equilibration were performed.

Homolytic breaks at the previously identified PYD short-arm C$\alpha$-C$\beta$ sacrificial bond ([56]) were introduced to the equilibrated structure using KIMMDY. Overall, 12 structures with different break patterns of four breaks each for the 16 PYD crosslinks were generated, taking care to include each residue three times and excluding crosslinks within 40 $\mathring{A}$ of each other or on the same triple helix. The radical-containing structures were equilibrated in the NpT ensemble for another 10 ns.

To determine suitable hyperparameters of the HAT GNN rate prediction (See above), three equilibrated collagen fibril systems with homolytic breaks are simulated for 500 ps, 5 ns and 50 ns with 50000 conformations per simulation. The pooled 5 ns and 50 ns data is used to determine the influence of number of conformations on the predicted reaction barrier (Fig. S6a). The distribution with 50000 and 100000 conformations have similar minium barriers of mean rate of 30.43 kcal/mol and 30.72 kcal/mol, respectively. The mean values are 56.08 kcal/mol and 55.33 kcal/mol. We conclude that the distributions with 50000 and 100000 conformations are similar enough to sample only 50000 conformations for further hyperparameter determination and the consecutive HAT simulations discussed in the main text. Using the same number of conformations, the impact of sampling for a longer time is limited (Fig. S6b,c). While the 50 ns sampling results in more different reactions, the lowest found barriers are similar for all sampling times. Looking at reactions with low hydrogen translation distances, *i.e.* those likely to have high reaction rates and therefore likely to occur, the MAE is between 4.29 kcal/mol and 5.07 kcal/mol for all three comparisons. This error is comparable to the error of the HAT GNN model (See above) when compared to QM calculations. Hence, a 500 ps sampling time is chosen for the collagen fibril simulations.

Predicting rates for all 50000 conformations would be prohibitively slow but the translation distance of the hydrogen atom can serve as a cheap heuristic to selectively predict on the conformations with the lowest barriers (Fig. S7a). Conformations that are not selected are assumed to have a rate of 0 1/s. When, for each reaction, only the 100 conformations with the lowest translation distance are evaluated, the sum of all evaluated rates is close to the sum of rates for all conformations for 90% of the reactions (Fig. S7b). 99% of the sum reaction rates are within $10^{-}2$ of the reference value for calculating the sum of rates with all conformations. However, not all sums of reaction rates are equally likely to be accurately determined by this scheme. For kMC, the relative rates determine the probability, *i.e.* the rate divided by the sum of all rates, of choosing a certain reaction. The selection probabilities for evaluating

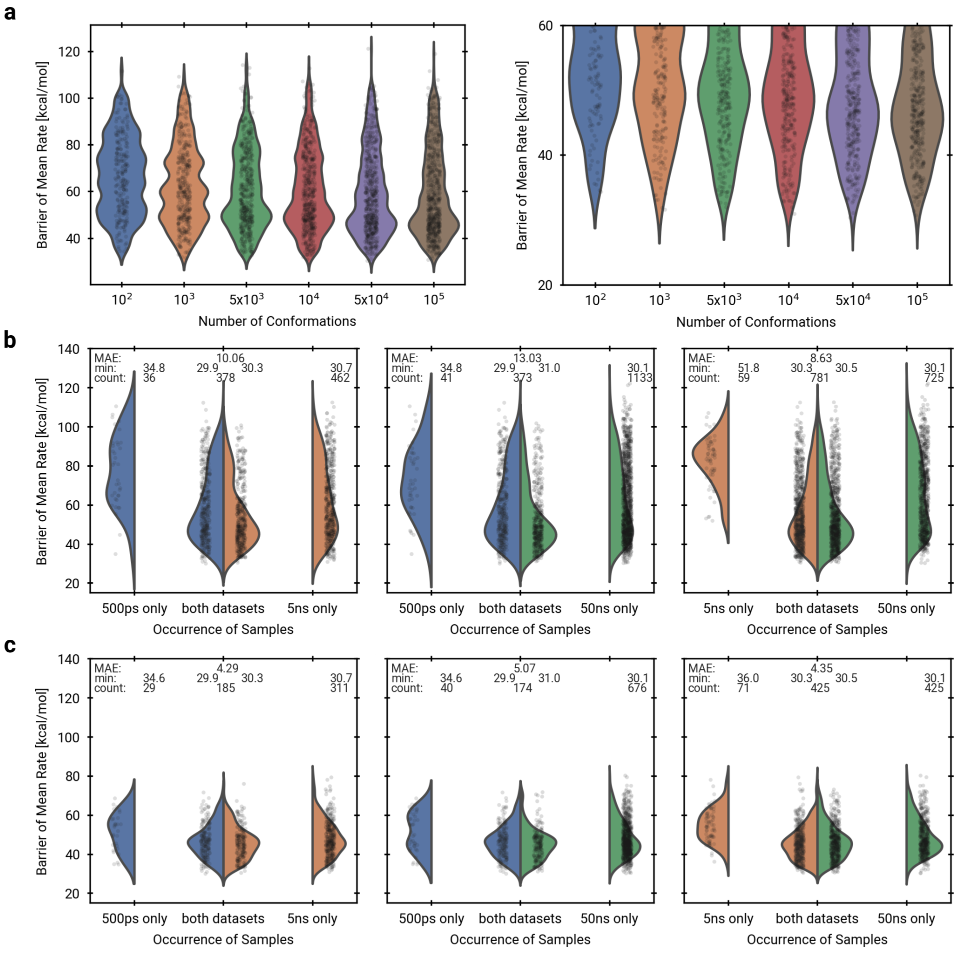

**Fig. S6 Hyperparameter tuning for the HAT GNN rate prediction of collagen fibril models: Number of conformations.** **a**, Relation between number of evaluated conformations and sampled barrier of mean rate for individual reactions. The dataset was generated from three simulations over 55 ns with a total of 100 000 conformations. For a smaller number of conformations every nth conformation is taken. As not all reactions were sampled in every conformation, the number of samples is distinct: n=230, n=370, n=476, n=517, n=581, n=612 (ordered by increasing number of conformations). The right panel is a cutout of the left panel. **b**, Comparison of the barrier of mean rate distributions for different sampling times of the same system. All predicted barriers are shown, as well as the minimum barrier in kcal/mol and number of samples for every set. In case a reaction occurs in both datasets, the MAE is shown in kcal/mol. **c**, Same as b, but only including reactions with at least one hydrogen translation distance below 2 Å.

all conformations (Fig. S7c) is similar to the selection probabilities for evaluating on only the 100 smallest translation distances (Fig. S7d). This can also be shown using the Brier divergence (Fig. S7e). For a single reaction, the barriers with the 100 smallest translations are shown in a time series (Fig. S7f). Overall, evaluating only the 100 conformations with the lowest hydrogen translation distance per reaction yields similar rates in this test compared to the full evaluation on 50000 conformations. The computational cost of the HAT GNN rate prediction is thus reduced by a factor of 500 for the collagen fibril model with this approach.

For state transitions after a reaction, the product was parametrised with the Grappa model '1.4.1-radical' (See above) and coordinates generated using the slow-growth method. HAT reactions were predicted for 100 structures with the smallest translation distance for each reaction with a overall prediction cutoff of 3 $\mathring{A}$. Rates were calculated using the Eyring equation assuming a transmission coefficient of 1.

The parameters for MD simulations were as follows: TIP3P water [57] was added to a box of 16.3 x 17.7 x 95.0 nm$^3$ and ions were NA$^+$ and Cl$^-$. During the NVT equilibration, the V-rescale [58] thermostat kept the temperature constant at 300 K with a coupling time of 0.1 ps. Protein heavy atoms were position restrained with a harmonic potential. The first 10 ns NpT equilibration at 1 bar used the Parrinello-Rahman barostat [59] with 2 ps coupling time and a 1 nN pulling force per chain to stretch the fibril. All subsequent equilibration and sampling simulations use the C-rescale [60] barostat with 5 ps coupling time and the aforementioned pulling force. The 500 ps sampling simulations write out protein coordinates every 10 fs for a total of 50000 frames analysed by the HAT plugin, whereas the 50 ns simulation writes out protein coordinates every 100 fs. Simulations for changing system coordinates to the product state using slow-growth are 10 ps long, have a 1ps coupling time for C-rescale and a 1fs time step. All other simulations have a time step of 2 fs with LINCS constraints [61] on h-bonds.

##### 1.7.3 Competing Reactions in Collagen

###### *system setup*

The solvated, neutralized and NpT-equilibrated structure of the collagen fibril from 1.7.2 was equilibrated under the different external forces ranging from 0 to 3 nN per chain for 2 ns and the convergence of the pulling simulation was checked based on the distances between the pull groups. The equilibration was followed by 2 ns of sampling simulations, during which bond distances where recorded with PLUMED [5]. Likewise, a single triple helix out of a collagen fibril and a single peptide out of the same triple helix where solvated in TIP3 water, neutralized with 0.15 M NaCl, energy minimized and equilibrated under NVT and NpT ensembles. The external forces applied range from no external force to 3 nN and in the case of the complete collagen fibril an additional, more physiological, sheared pulling was introduced by sampling the force on each triple helix from a gaussian distribution with a standard deviation of 30 % of the mean force centred around the nominal external force.

###### *Hydrolysis reaction rates*

The force on each bond is calculated based on the morse potential and extension of each bond according to Eq. 6.

$$F = 2\ \beta\ E_{dis}\ e^{-\beta\ \Delta r}\ (1 - e^{-\beta\ \Delta r}) \tag{6}$$

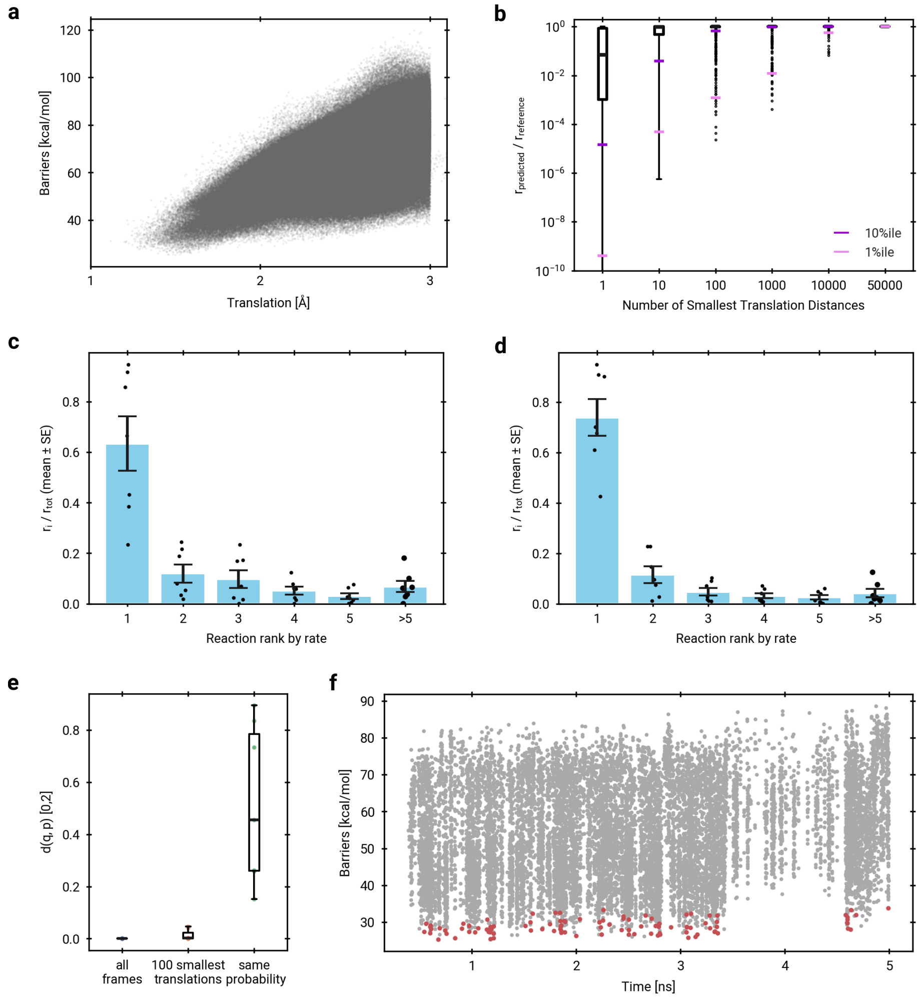

**Fig. S7 Hyperparameter tuning for the HAT GNN rate prediction of collagen fibril models: Number of predictions. a**, Relation between translation distance of the hydrogen atom for the HAT reaction and the barrier of that reaction. From a single simulation of 5 ns length with 50000 conformations, 963518 reaction predictions for different reactions are shown. **b**, Boxplot of the fraction of the sum of all rates per reaction calculated with a subset of all predictions. For each reaction, only the n conformations with the smallest translation distance are chosen and evaluated (n=1084 reactions). **c**, Selection probability of reactions by calculating the sum of all rates for all conformations. For 9 simulations of 5 ns length with 500000 conformations, reactions are ranked by their rate. The contribution of the rates to the sum of all rates per simulation is averaged over the ranks. Bars indicate the mean, error bars the standard error of the mean. **d**, Selection probability of reactions by calculating the sum of all rates for conformations with the 100 smallest translation distances per reaction. The data is the same as in c. **e**, Brier divergence compared to a uniform rate distribution. Same data as in c. **f**, Barriers over time for a single reaction chosen for its low barrier. The 100 conformations with the lowest translation distance are shown in red, all others are shown in grey.

with $\beta = \sqrt{k/(2\ E_{dis})}$,the spring constant $k$, and the dissociation energy $E_{dis}$.

To account for the chemical environment of each bond as well as entropic effects of the simulation setup and compare rates directly to experimental reference data, the average force per bond at 0 nN external force is subtracted from the average force. Then the force can be translated to a reaction rate either directly based on single molecule force spectroscopy results by [62] or calculated from reaction barriers from QM calculations from the same publication. The resulting force-response curve and sampled reaction rates can be seen in Fig. A4b and c respectively, using two smaller systems (rendered in d), a single peptide and a triple helix, both excerpts from the large collagen fibril.

On top of the Solvent Accessible Surface Area (SASA) of the peptide bond, the hydrolysis plugin can take into account further parameters such as the pH value, which manifests as a modified attempt frequency based on the concentration of $OH^-$ (i.e. the probability of any given attacking water molecule being deprotonated) due to the reaction being base-catalysed. Fig. A4e) shows this for the single peptide and triple helix systems.

The more basic the pH value the higher the reaction rate as a result of more successful attempts. When it comes to selecting the water molecule for the topology modification to execute the chosen reaction, additional factors are considered such as the distance and angle of attack according to the Bürgi-Dunitz angle [63]. Future iterations of the hydrolysis plugin could incorporate this angle into the rate calculation as well. The code for the kimmdy-hydrolysis plugin is available at https://github.com/graeter-group/kimmdy-hydrolysis.

##### 1.7.4 DNA simulations

###### *DNA sequences*

We provide the DNA sequences for each sample in Fig. S8. All sequences are designed such that additional thymine-thymine sites are avoided.

###### *Parametrisation of the dimer residue*

To perform consecutive reactions a parametrisation of the dimer residue is necessary since it does not appear in the conventional DNA force field. To achieve this we create a modified thymine residue in the OL21 force field [64].

We obtain a Hartree–Fock optimized structure of the thymine–thymine dimer (with no backbone connectivity between the thymines). From this structure we calculate partial charges *via* the Restrained Electrostatic Potential (RESP) fitting method [65] utilizing AmberTools [7], where we constrain the partial charges of the sugar to the values already present in the force field. The resulting partial charges are given in table S5.

Through the reaction four $sp^2$-hybridized carbon atoms change to $sp^3$ hybridisation. To cover this we change the atom types in the following way:

$$\mathrm{C5} : \mathrm{CM} \rightarrow \mathrm{CT}$$
$$\mathrm{C6} : \mathrm{CM} \rightarrow \mathrm{CT}$$

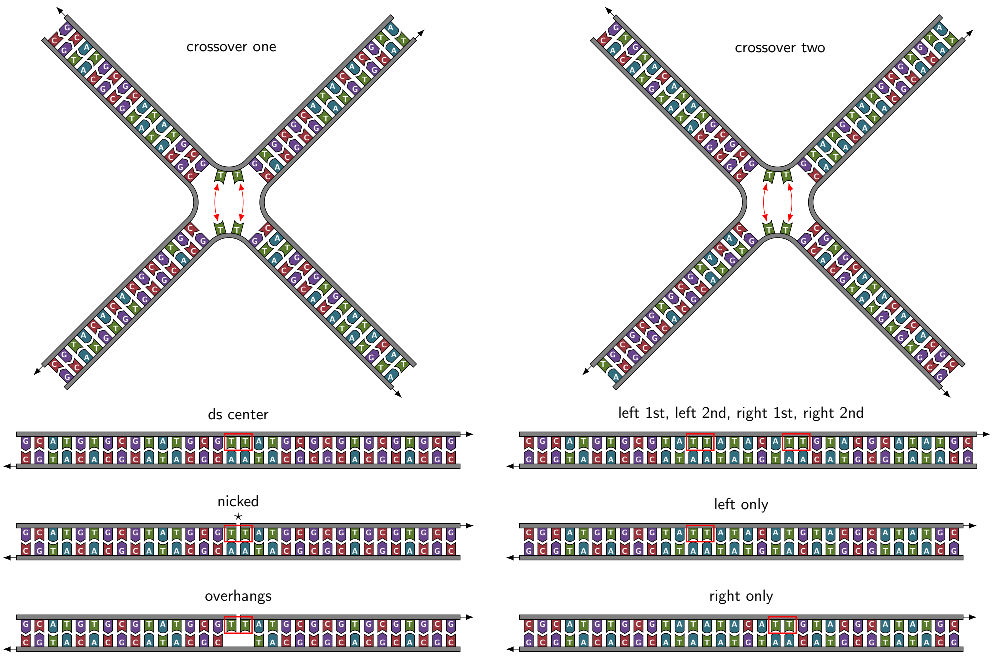

**Fig. S8 DNA sequences used in simulations.** Schematic representations of DNA structures used in simulations providing the entire nucleotide sequence of each sample. Red arrows and red boxes denote thymines considered for dimerisation by KIMMDY. Black arrows denote 3'-ends.

**Table S5** Partial atomic charges for atoms in a thymine residue being part of a CPD unit. Sugar or phosphate atoms are not shown, since they have not been modified.

| Atom name | Partial charge [e] |
|---|---|
| N1 | -0.36457 |
| C6 | 0.01612 |
| H6 | 0.21094 |
| C5 | -0.09999 |
| C7 | -0.44844 |
| H71 | 0.13809 |
| H72 | 0.13809 |
| H73 | 0.13809 |
| C4 | 0.93659 |
| O4 | -0.62419 |
| N3 | -0.78479 |
| H3 | 0.39374 |
| C2 | 0.85606 |
| O2 | -0.63253 |

$$H6 : H4 \rightarrow H1$$

$$N1 : N^* \rightarrow N$$

Additionally, the improper dihedrals C4–C6–C5–C7 and N1–C5–C6–H6 are removed in comparison to the unmodified thymine residue. After changing the atom type, bonded angle interactions are added to the force field as educated guesses based on parameters present in the force field for similar atom type combinations (see table S6).

**Table S6** Angle interactions added to the forcefield to accommodate the changed atom types.

| Atom 1 | Atom 2 | Atom 3 | Function type | Equilibrium angle [°] | Force constant [kJ·mol$^{-1}$·rad$^{-2}$] |
|---|---|---|---|---|---|
| OS | CT | N | 1 | 109.5 | 418.40 |
| CT | C | NA | 1 | 114.1 | 585.76 |
| N | C | NA | 1 | 115.4 | 585.76 |
| H2 | CT | N | 1 | 109.5 | 418.40 |

##### *MD simulations*

Initial topologies of all DNA systems—including the complex systems from the main text as well as the benchmark systems TdT and $dT_{20}$ —were built using OxView [66]. The crossover structures were pre-equilibrated in the coarse-grained representation using OxView's built-in dynamics module. Structures were then converted to all-atom PDB files via TacoxDNA [67]. All systems were solvated with TIP3P water (in cubic boxes with edge lengths of 5 nm for TdT, 10 nm for dT20, 18 nm for crossover, and 15 nm for all other systems) and neutralized with $Na^+$ ions using GROMACS. Energy minimization was performed on all structures using the steepest descent algorithm. All MD simulations were performed with the OL21 nucleic forcefield [64] extended by the custom dimer residue.

For the benchmark systems TdT and $dT_{20}$, a 5 ns NVT simulation was followed by a 5 ns NPT simulation. Subsequently, 100 ns of MD simulations were performed without the use of KIMMDY, with three independent runs per system, all starting from the same initial PDB file. The TdT system was then simulated using KIMMDY, consisting of a 5 ns MD simulation followed by a 100 ns MD simulation, during which reaction rates were sampled every 1 ps. Three independent KIMMDY runs were performed, each starting from one of the previously obtained NPT configurations.

For the systems ds center, nicked, overhangs, left only, right only, crossover one, crossover two (jointly referred to as crossover in the main text), left 1st, and right 1st, a 5 ns NVT simulation was followed by a 5 ns NPT simulation. KIMMDY simulations were then performed, consisting of a 10 ns MD simulation followed by a 100 ns production run, with reaction rates sampled every 1 ps. Each system was simulated three times independently, with all runs sharing the same initial PDB file. Note that rates for left 1st and right 1st were obtained from the same simulations, as neither site had reacted at that point.

For the left 2nd and right 2nd systems, a total of nine KIMMDY runs were conducted. These runs started from one of the NPT configurations previously obtained for the left 1st/right 1st system. Each run consisted of a 5 ns MD simulation, followed by a second 5 ns MD simulation during which the first reaction event was identified. After a brief relaxation, a 100 ns MD simulation was performed, with KIMMDY sampling reaction rates every 1 ps. Of the nine runs, five resulted in reactions at left 2nd, and four at right 2nd.

##### *Model for dimerisation*

Reaction rates for thymine-thymine dimerisation have been modelled according to Eq. 7, taking into account the distance $d$ between the $C_5 = C_6$ double bonds as well as the $C_5^a - C_6^a - C_6^b - C_5^b$ (for thymines a and b) dihedral angle $\eta$.

$$r = 1\ \mathrm{ps}^{-1} \cdot \exp\left(-(k_1 \cdot |d - d_0| + k_2 \cdot |\eta - \eta_0|)\right) \tag{7}$$

The 1 $\mathrm{ps}^{-1}$ time scale is taken since dimerisation in a favourable configuration is reported to be an ultrafast photoreaction that occurs in less than 1 ps [68]. $k_1$ and $k_2$ are parameters denoting the effect of deviation from the optimal configuration for distance and angle deviations respectively. The distances $d_0$ and $\eta_0$ have been obtained from a geometry optimized (Hartree-Fock level) structure of the CPD product, with $d_0 = 0.157$ nm and $\eta_0 = 16.744°$.

From this, an attempted reaction is considered to be successful and contributing to the quantum yield $\phi$ if the reaction rate is greater than the rate of the vibrational cooling (VC), which is reported to be 0.5 $\mathrm{ps}^{-1}$ [69]:

$$\phi = \frac{\#\ \text{frames with}\ r > 0.5\ \mathrm{ps}^{-1}}{\#\ \text{frames}} \tag{8}$$

For systems with multiple adjacent thymine-thymine-dimerisation sites, the normalization is performed per thymine (equation 9).

$$\phi = \frac{2}{N_T} \cdot \frac{\#\text{TT-pairs with}\ r > 0.5\ \mathrm{ps}^{-1}}{\#\text{frames}} \tag{9}$$

To obtain values for $k_1$ and $k_2$ we considered two well-described benchmark structures, the thymine-thymine dinucleotide TdT and the single-stranded polyT DNA $dT_{20}$. The quantum yield is reported to be 0.013 for TdT and 0.028 for $dT_{20}$[70]. Using these values, a grid search has been performed to screen possible values for $k_1$ and $k_2$. For each system, three 100 ns MD runs (see below for MD details) have been considered, writing out the coordinates every 4 ps, yielding a total of 75000 configurations per system. For TdT the predicted quantum yield for each $k_1, k_2$ combination was determined from Eq. 7 and 8, for $dT_{20}$ Eq. 7 and 9 were used. The results of the grid searches are shown in Fig. S9a and S9b.

Combining the results of both grid searches by using the maximum of the deviation from the respective experimental quantum yield and selecting the $k_1, k_2$ combination with the lowest deviation from either experimental quantum yield provides a set of parameters that simultaneously covers both experimental values for the benchmark

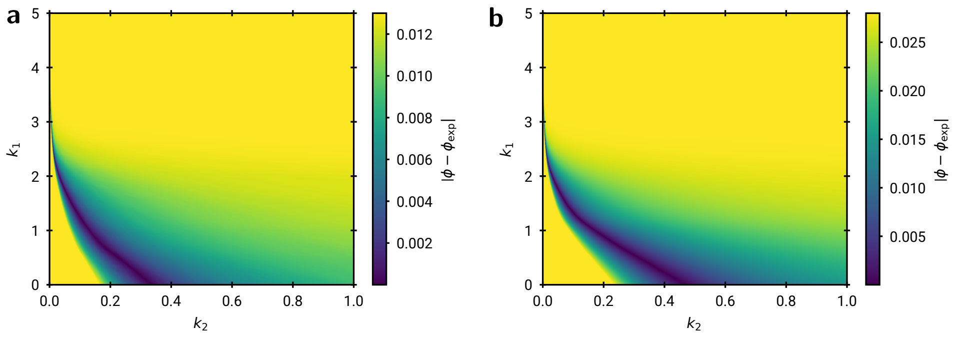

**Fig. S9 Grid search results for TdT and dT$_{20}$.** **a**, Absolute deviation from experimental quantum yield for different values of $k_1, k_2$ for the TdT system. Deviations have been capped at the value of the experimental quantum yield (0.013). Based on configurations from three independent 100 ns MD simulations. **b**, Absolute deviation from experimental quantum yield for different values of $k_1, k_2$ for the dT$_{20}$ system. Deviations have been capped at the value of the experimental quantum yield (0.028). Based on configurations from three independent 100 ns MD simulations.

systems. The parameters used for the remainder of the study are given by

$$k_1 = 2.017, \quad k_2 = 0.030, \quad \text{max. deviation from } \phi_{\text{exp}} \approx 8 \cdot 10^{-5}.$$

To verify this selection, we calculate the rate for threshold structures from Law *et al.* [71] ($d = 0.370$ nm, $\eta = 48.2°$), Johnson *et al.* [72] ($d = 0.340$ nm, $\eta = 27.0°$) and McCullagh *et al.*[73] ($d = 0.352$ nm, $\eta = 40.0°$ [no angle was given, therefore arbitrarily set to 40.0°]) shown in table S7. This evaluation shows that the selected parameters yield reasonable values that reach our reaction threshold of 0.5 ps$^{-1}$ for at least one of the previously reported threshold structures.

**Table S7** Reaction rates for different threshold structures calculated with Equation 7 using the selected $k_1, k_2$ combination.

| Threshold | $r$ [ps$^{-1}$] |
|---|---|
| $d = 0.370$ nm, $\eta = 48.2°$ | 0.253 |
| $d = 0.340$ nm, $\eta = 27.0°$ | 0.508 |
| $d = 0.352$ nm, $\eta = 40.0°$ | 0.336 |

The implementation can be found at https://github.com/graeter-group/kimmdy-dimerization. This implementation does not include other possible photoproducts such as the 6-4 photoproduct or *cis-anti* and *trans-anti* isomers. (see Fig. S10).

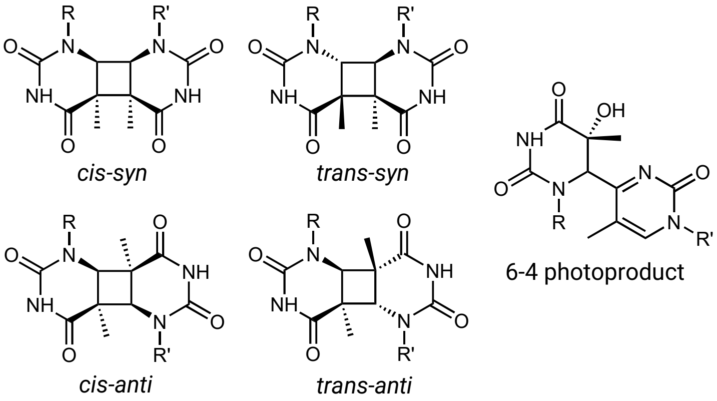

**Fig. S10 Possible thymine-thymine-photodimerisation products.** Only the *cis-syn* and the *trans-syn* products are covered by the presented KIMMDY plugin. For R ≠ R' additional enantiomers for the *cis-syn* and *trans-anti* products exist. The shown *trans-syn* and *cis-anti* products always have an additional enatiomer, even if R = R'.

### 2 Supplementary Notes

#### 2.1 Kinetic Monte Carlo

##### 2.1.1 Introduction

Monte Carlo (MC) methods cover approaches where random objects or processes are generated computationally and are used in a wide range of areas such as industrial engineering, materials science and finance [74] . In many applications, the "randomness" is introduced artificially to estimate a deterministic quantity, such as the computation of a finite integral over many dimensions. Previous works have combined MD with MC, such as ref. [75] , where the authors created a hybrid molecular dynamics/Monte Carlo algorithm for the atomistic simulations of precipitation in alloys.

In the present paper, reactions are considered to be rare and therefore happen on time scales which are much longer than typically accessible through molecular dynamics. Therefore, such reactions will not happen fast enough to be observable in direct simulations, even if topological changes through bond breakage/formations were allowed in parametrized classical force fields. The typical kinetic energy of the particles would be insufficient to overcome a reaction barrier during a simulation run, and a transition of the system state would not be observable.

Kinetic Monte Carlo (kMC) is a type of MC method that makes it possible to access much larger time scales than MD typically can, by treating each reaction as a random process with a fixed mean rate of occurrence. Once an event has been appropriately sampled, the associated reaction or change is performed, and the simulation time is adjusted. This means that kMC can stochastically simulate the behaviour of rare events, which would be typically too slow for MD simulations. One of the first applications was a kMC algorithm for simulating an Ising spin model [**?** ] by Bortz et al. Variations of the kMC algorithm have been applied under various names, but the terminology has settled on kinetic Monte Carlo only in the 1990s [76].

Typically, a fixed table of rates is precomputed for all possible reactions. However, kMC methods with on-the-fly computed rates have also been developed. Henkelman and Jónsson [**?** ] applied harmonic transition state theory in combination with the dimer transition state search method [77] to dynamically find possible reactions and compute their respective reaction rates.

##### 2.1.2 Theoretical background

We here show the well-known statistical background which underpins the kMC algorithm. The interested reader is referred to [78] for further reading. Similarly, we refer the reader to [**?** ] for a general introduction to Poisson point processes.

###### *Single process*

Consider some random process of which we only know that an event occurs on average $M$ times within an observation time $t_{obs}$, i.e. at an average rate or intensity of $\lambda = M/t_{obs}$ per unit time. Furthermore, we expect the events to occur at random and independently of each other.

We can take some interval $[0, t)$ on the timeline and subdivide this into $n$ smaller time increments of $\Delta t$. The number of events in the total interval can be described by a Binomial distribution with success probability $p = \lambda \Delta t$, and therefore with an average of $np = (t/\Delta t)\lambda \Delta t = \lambda t$ events occurring. As $\Delta t \to 0$, the Binomial distribution becomes a Poisson distribution with mean $\lambda t$. This type of process can be modelled as a (homogeneous) Poisson point process (PPP).

The probability that exactly 0 events occur in time $t$ is given by the Poisson distribution,

$$p_\lambda^0(t) := \mathbb{P}(k = 0; \mu = \lambda t) = \frac{e^{-\mu}\mu^k}{k!} = e^{-\lambda t}, \tag{10}$$

where $k$ is the number of event occurrences and $\mu$ is the distribution mean parameter.

Consequently, the probability that the time of first occurrence $T$ is greater than $t$ is given by

$$\mathbb{P}(T > t) = e^{-\lambda t}. \tag{11}$$

Taking the complement (first occurrence within time $t$) gives

$$F_T(t) = \mathbb{P}(T \leq t) = 1 - e^{-\lambda t}, \tag{12}$$

which is the cumulative distribution function (CDF) of the exponential distribution. Therefore the time of first occurrence is an exponentially distributed random variable with mean parameter $\lambda$.

Let us turn to the situation in which we can produce an event from a PPP, but then need to adjust the time that has passed until this event occurred. The process occurs randomly, therefore we cannot say for certain how much time has passed. Instead, it must be consistent with the probability density function (PDF) of first occurrence and therefore needs to be appropriately sampled.

This can be accomplished using inverse transform sampling (cite), resulting in a time step of

$$t_{\text{step}} = \log{(1/u)}/\lambda, \tag{13}$$

where $u$ is a number sampled from the standard uniform distribution. The time of first occurrence can therefore be sampled using Eq. 13.

***Multiple competing processes***

Let there be $N$ independently competing Poisson point processes (PPP), each of which is indexed by $j$ with average rate $\lambda_j$. Which process will occur first? Each process will have a different PDF for the first occurrence.

The time of first occurrence over all processes $T$ is given by

$$T = \min{(T_1, \ldots, T_N)}, \tag{14}$$

where $T_j$ is the time of first occurrence random variable of process $j$. The probability that $T$ is larger than some $t$ is given by

$$\mathbb{P}(T > t) = \mathbb{P}(T_1 > t, \ldots, T_N > t) \tag{15}$$

$$= \prod_j^N \mathbb{P}(T_j > t) = \prod_j^N (1 - \mathbb{P}(T_j \leq t)) \tag{16}$$

$$= \prod_j^N e^{-\lambda_j t} = e^{-(\sum_j^N \lambda_j)t}, \tag{17}$$

where we have used the independence property of the PPPs. Therefore the CDF of $T$ is given by Eq. 12 with $\lambda = \sum_j^N \lambda_j$.

$$\mathbb{P}(T \leq t) = 1 - e^{-(\sum_j^N \lambda_j)t}, \tag{18}$$

which is equivalent to a random variable with rate $\left(\sum_j^N \lambda_j\right)$. The the reaction time of the competing processes can therefore be modelled as a single PPP with a rate equivalent to the summed rates and sampled using Eq. 13.

The probability that an event from process $J$ occurred first is given by

$$\mathbb{P}(\forall_{j \neq J} \; J \leq T_j) = \int_0^\infty \left. \frac{F_T(v)}{dv} \right|_t \mathbb{P}(\forall_{j \neq J} \, T_j > t) dt \tag{19}$$

$$= \int_0^\infty \lambda_J e^{-\lambda_J t} \prod_{j \neq J}^N e^{-\lambda_j t} dt = \lambda_J \int_0^\infty e^{-(\sum_j^N \lambda_j)t} dt \tag{20}$$

$$= \frac{\lambda_J}{\sum_j^N \lambda_j}. \tag{21}$$

***Poisson point process with non constant rate***

When the mean rate of the Poisson point process $\lambda$ is not constant, but changes with time this becomes a non-homogeneous Poisson point process.

We consider the case of discrete time steps $\Delta s$, where the mean rate $\lambda_s$ changes with every time index $s$. Let $\lambda_s$ denote the mean rate from time $s$ up to $s + \Delta s$.

Consider a finite time $S$ that is subdivided into $n$ intervals of equal length, s.t. $n = S/\Delta s$ and indexed by $s$. Further, let each interval $[s, s+\Delta s]$ have a separate PPP with mean rate $\lambda_s$.

The probability that no event occurs until time $S$ in every interval is then given by

$$\mathbb{P}(\forall_s \, T_s > \Delta s) = \prod_s^n e^{-\lambda_s \Delta s} = \prod_s^n e^{-\lambda_s S/n} \tag{22}$$

$$= e^{-(\sum_s^n \lambda_s/n)S}, \tag{23}$$

where $T_s$ is the time of first occurrence in interval $[s, s + \Delta s]$. Therefore, this joint process is defined by the survival function of an exponentially distributed random variable with mean rate $\sum_s^n \lambda_s/n$. This is equivalent to a single PPP with an averaged mean rate over all intervals. In practise this assumes that the observation time of the changing rates is large enough such that the empirical mean is close enough to the true mean of the rates.

##### 2.1.3 The kinetic Monte Carlo algorithm

The kinetic Monte Carlo (kMC) algorithm can be readily constructed using the results of section 2.1.2.

The setup is $N$ independently competing reactions that are modelled as Poisson point processes with rates $\lambda_i(t)$, at simulation time $t$.

Over the entire MD simulation run of time $T$, which is subdivided into $n$ time steps of length $\Delta t$, each reaction has an average mean rate of $r_i = \sum_{j=1}^n \lambda_i(t_j)/n = (\Delta t/T)\sum_{j=1}^n \lambda_i(t_j)$.

The selection probability of each reaction is given by Eq. 21,

$$p_i = \frac{r_i}{\sum_{j=1}^N r_j}. \tag{24}$$

The time is then progressed according to Eq. 13,

$$t_{\text{new}} \to t_{\text{old}} + \frac{\log(1/u)}{\sum_{j=1}^N r_j}, \tag{25}$$

where $u$ is sampled from a standard uniform distribution.

Our approach is a variation of adaptive kMC [79], because the event list of independently competing reactions is generated on-the-fly for each visited state.

#### 2.2 Calculating reaction rates from barriers

Arrhenius described an empirical relation [80] between activation energy $E_a$ and rate constant using a pre-exponential factor $A$ and the gas constant $R$ as well as the temperature $T$:

$$k = A\exp\left(\frac{-E_a}{RT}\right) \tag{26}$$

Following a statistical mechanics perspective, Eyring found an expression for $A$ [81] that forms the basis for transition state theory (TST):

$$\begin{aligned} A &= \kappa_0 \frac{k_B T}{h} \\ k_{\text{TST}} &= \kappa_0 \frac{k_B T}{h}\exp\left(\frac{-E_a}{RT}\right) \end{aligned} \tag{27}$$

Here, $k_B$ is the Boltzmann constant, $h$ is the Planck constant and $\kappa_0$ is the transmission coefficient.

Several extensions to TST exist that use different expression for the transmission coefficient $\kappa_0$, including harmonic TST, where vibrational modes at the reactant and transition state are calculated to estimate the likelihood of barrier crossing events [82]. In practice, $\kappa_0$ is often estimated to be in the range of 1 - 10 [83].

This simplifies the task of finding a rate constant to finding a reaction barrier. Many QM-based methods exist to find transition states from energy minima, including the nudged elastic band method and gradient based search methods like the Dimer method [77, 84]. In this work, the QM transition state search is emulated with a combined MD/ML approach.

[22] Chan, B., Collins, E. & Raghavachari, K. Applications of isodesmic-type reactions for computational thermochemistry **11**, e1501. URL https://onlinelibrary.wiley.com/doi/abs/10.1002/wcms.1501. _eprint: https://onlinelibrary.wiley.com/doi/pdf/10.1002/wcms.1501.

[23] Hehre, W. J., Ditchfield, R., Radom, L. & Pople, J. A. Molecular orbital theory of the electronic structure of organic compounds. v. molecular theory of bond separation **92**, 4796–4801. URL https://doi.org/10.1021/ja00719a006. Publisher: American Chemical Society.

[24] Luo, Y.-R. *Comprehensive Handbook of Chemical Bond Energies* (CRC Press).

[25] Neese, F. The ORCA program system **2**, 73–78. URL https://wires.onlinelibrary.wiley.com/doi/10.1002/wcms.81. Publisher: John Wiley & Sons, Ltd.

[26] Neese, F. Software update: The ORCA program system—version 5.0 **12**, e1606. URL https://onlinelibrary.wiley.com/doi/abs/10.1002/wcms.1606. _eprint: https://onlinelibrary.wiley.com/doi/pdf/10.1002/wcms.1606.

[27] Zhao, Y. & Truhlar, D. G. The m06 suite of density functionals for main group thermochemistry, thermochemical kinetics, noncovalent interactions, excited states, and transition elements: two new functionals and systematic testing of four m06-class functionals and 12 other functionals **120**, 215–241. URL https://doi.org/10.1007/s00214-007-0310-x.

[28] Treyde, W., Riedmiller, K. & Gräter, F. Bond dissociation energies of X–H bonds in proteins. *RSC Advances* **12**, 34557–34564 (2022). URL https://xlink.rsc.org/?DOI=D2RA04002F.

[29] Neese, F. in *First principles approach to spin-hamiltonian parameters* 1 edn, (ed.Misra, S. K.) *Multifrequency Electron Paramagnetic Resonance* 295–326 (Wiley). URL https://onlinelibrary.wiley.com/doi/10.1002/9783527633531.ch6.

[30] Stoll, S. & Schweiger, A. EasySpin, a comprehensive software package for spectral simulation and analysis in EPR **178**, 42–55. URL https://www.sciencedirect.com/science/article/pii/S1090780705002892.

[39] Tsang, W., Awan, I. A., McGivern, W. S. & Manion, J. A. Soot precursors from real fuels: The unimolecular reactions of fuel radicals .

[40] Awan, I. A., Burgess, D. R. & Manion, J. A. Pressure Dependence and Branching Ratios in the Decomposition of 1-Pentyl Radicals: Shock Tube Experiments and Master Equation Modeling. *The Journal of Physical Chemistry A* **116**, 2895–2910 (2012).

[41] Comandini, A., Awan, I. A. & Manion, J. A. Thermal decomposition of 1-pentyl radicals at high pressures and temperatures. *Chemical Physics Letters* **552**, 20–26 (2012).

[42] Yamauchi, N., Miyoshi, A., Kosaka, K., Koshi, M. & Matsui, H. Thermal Decomposition and Isomerization Processes of Alkyl Radicals. *The Journal of Physical Chemistry A* **103**, 2723–2733 (1999).

[43] Tsang, W., Walker, J. A. & Manion, J. A. The decomposition of normal hexyl radicals. *Proceedings of the Combustion Institute* **31**, 141–148 (2007).

[44] Tsang, W., Awan, I. A., McGivern, W. S. & Manion, J. A. Soot precursors from real fuels: The unimolecular reactions of fuel radicals .

[80] Arrhenius, S. Über die Reaktionsgeschwindigkeit bei der Inversion von Rohrzucker durch Säuren. *Zeitschrift für Physikalische Chemie* **4U**, 226–248 (1889). URL https://www.degruyter.com/document/doi/10.1515/zpch-1889-0416/html.

[81] Eyring, H. The Activated Complex in Chemical Reactions. *The Journal of Chemical Physics* **3**, 107–115 (1935). URL https://pubs.aip.org/jcp/article/3/2/107/203352/The-Activated-Complex-in-Chemical-Reactions.

[82] Truhlar, D. G., Garrett, B. C. & Klippenstein, S. J. Current Status of Transition-State Theory (1996).

[83] Andersen, M., Panosetti, C. & Reuter, K. A Practical Guide to Surface Kinetic Monte Carlo Simulations. *Frontiers in Chemistry* **7** (2019). URL https://www.frontiersin.org/articles/10.3389/fchem.2019.00202.

[84] Henkelman, G., Uberuaga, B. P. & Jónsson, H. A climbing image nudged elastic band method for finding saddle points and minimum energy paths. *The Journal of Chemical Physics* **113**, 9901–9904 (2000). URL https://pubs.aip.org/jcp/article/113/22/9901/185050/A-climbing-image-nudged-elastic-band-method-for.
